# A 28-year evolution experiment on *Burkholderia pseudomallei* survival in nutrient-depleted sterile water

**DOI:** 10.64898/2026.02.06.704356

**Authors:** Chalita Chomkatekaew, Premjit Amornchai, Sayan Langlah, Janjira Thaipadungpanit, Elizabeth M Batty, Direk Limmathurotsakul, Kalayanee Chairat, Kanyarat Sueksakrit, Siribha Apinan, Phornpimon Tipthara, Joel Tarning, Víctor Rodríguez-Bouza, John A. Lees, Jukka Corander, Nicholas PJ Day, Nicholas J White, Nicholas R Thomson, Vanaporn Wuthiekanun, Julian Parkhill, Claire Chewapreecha

**Author notes:** Correspondence to: Claire Chewapreecha. Contributed equally (Vanaporn Wuthiekanun, Julian Parkhill, Claire Chewapreecha). Deceased, 1 February 2026 (Nicholas J White).

## Abstract

Environmental persistence allows opportunistic pathogens to survive a range of harsh conditions, increasing the likelihood of eventual infection. *Burkholderia pseudomallei*, the causative agent of melioidosis, can endure long-term nutrient-depletion in the environment, but its adaptive mechanisms remain poorly understood. Here, we investigated the evolutionary trajectory of a clinical *B. pseudomallei* strain maintained in sterile water since 1994. The strain was inoculated into nine individual tubes at nine initial concentrations (10^2^-10^10^ CFU/mL) and has remained viable to date. Liquid chromatography-mass spectrometry analysis of the water identified potential carbon sources, including phthalic acid — a plastic degradation product likely leached from inoculation tubes — which the strain can metabolise via an intact catabolic operon. Genomic variations accumulated between 1994 and 2022 were characterised using both single-colony and plate-sweep sequencing which provided complementary insights. Across all tubes, we identified 249 single-nucleotide polymorphisms (SNPs), 73 indels, and a large-scale deletion. Cultures from each tube displayed a consistently low mutation rate (3.18 × 10⁻⁸ SNPs per site per year), suggesting that cells entered a dormant or slow-growth state. Of 393 genes with mutations, 193 were independently mutated in more than one tube, particularly those involved in signal transduction, cell wall and membrane biogenesis, and secondary metabolite synthesis. These patterns indicate parallel adaptation to long-term nutrient deprivation through modulation of cell-density–related functions and loss of metabolically costly pathways. Remarkably, *B. pseudomallei* from this experiment remain viable after nearly three decades, providing a rare natural model for understanding how environmental bacteria endure and adapt in extremely nutrient-depleted conditions.

## Introduction

*Burkholderia pseudomallei* is a Gram-negative environmental bacterium found in soil and water across tropical and subtropical regions, causing melioidosis outbreaks in northern Australia^1^, South-and Southeast Asia^2,3^, parts of Africa^4^, and more recently in the Americas^5^. Humans become infected through environmental exposure – with bacterial acquisition mediated by percutaneous inoculation, inhalation, or ingestion – leading to melioidosis, which can present as acute disease (85-90% of cases), chronic disease with persistent symptoms (9-12%), or latent, asymptomatic infection (<5%) that can reactivate years later^6,7,1^. Both chronic and latent infections reflect the bacterium’s remarkable capacity for long-term persistence within the host, mediated by slow replication or entry into a dormant state. A frequently cited example of latent infection is the so-called “Vietnamese time bomb”, describing a case in which a US veteran developed melioidosis years after military service in Vietnam, in the absence of any identified subsequent exposure^8^. In the environment, *B. pseudomallei* survives harsh conditions, including desiccation^9^, low pH ^10^, severe climates^11–14^, intracellular life within other organisms such as amoeba^15^, and prolonged nutrient depletion^16,10,17–22^. This environmental persistence is a prerequisite for infection, facilitating both pathogen survival outside the host and subsequent disease acquisition.

The bacterium’s persistence and adaptation may be supported by its large and flexible genetic repertoire, consisting of two stable chromosomes (total size approximately 7.24 Mbp)^23^. It has an open pan-genome containing an approximate 5,059 core genes (range 4,067-5,905) and around 8,920 accessory genes (range 1,075-21,748)^24–26,22^. Within individual genomes, multiple gene paralogues diverge in sequence and potential function, allowing for diversification of biological function. At the population level, each gene — including its paralogues – can exist as multiple alleles, providing the capacity for interindividual adaptation. An allele-based study of global *B. pseudomallei* populations has identified loci under co-selection with multiple genes, suggesting these loci may act as evolutionary hubs. Notably, some of these genes are expressed under nutrient starvation^21^, indicating a role in adaptation to environmental stress. Population-scale genomic analysis from Thailand, a disease hotspot, further revealed that lineage-specific accessory genes may facilitate survival under environmental pressures, including nutrient depletion^22^. These genomic observations are supported by field evidence. In northeast Thailand and other endemic areas, *B. pseudomallei* is consistently detected more frequently in nutrient-poor rice paddies compared to nutrient-rich ones^20,27^, suggesting an ecological advantage in oligotrophic environments. However, despite population-level patterns and candidate genes being identified, the mechanisms and evolutionary pathways underlying long-term adaptation to nutrient limitation remain largely unexplored.

To address this, we leveraged a unique long-term *in vitro* experiment initiated in 1994 by Mrs Vanaporn Wuthiekanun and Professor Nicholas J White. In this study^16,18^, a clinical strain of *B. pseudomallei* was inoculated into sterile, triple-filtered distilled water and stored in semi-closed 10 mL plastic tubes (used instead of glass due to BSL-3 safety regulations) at ambient temperature. Nine different starting inoculum concentrations were used ranging from 10^2^ to 10^10^ CFU/mL (**Figure 1a**, **Supplementary Table 1**). Bacterial viability has been assessed regularly by plating, and remarkably, viable bacteria have persisted in all tubes for nearly 30 years. This rare, time-resolved, and controlled system offers an unprecedented opportunity to investigate both short-and long-term genomic adaptations to nutrient deprivation. The findings offer insights into how *B. pseudomallei* persists in the environment and may inform understanding of long-term chronic and latent infections in humans.

**Figure 1.**
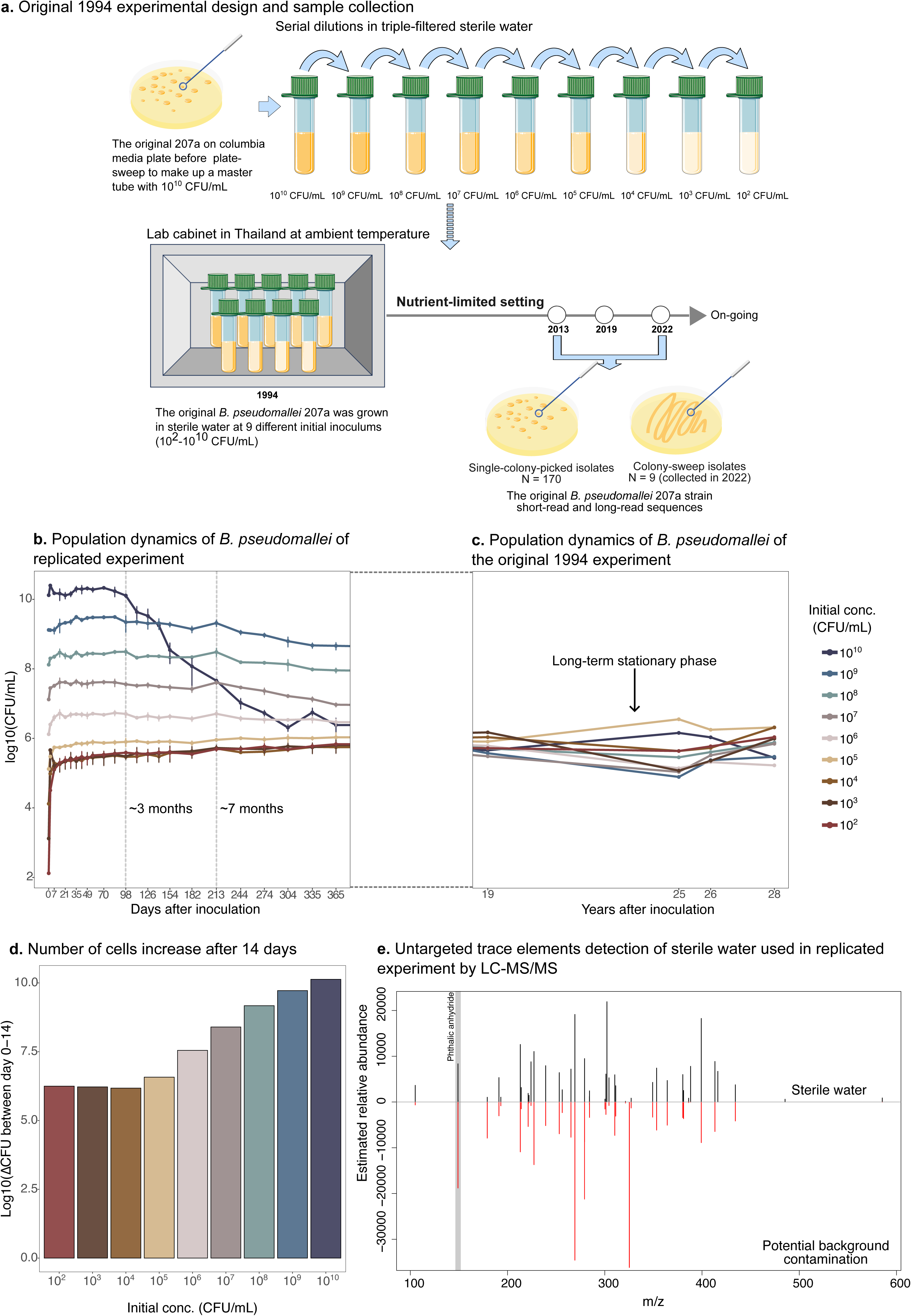
Experimental design, population dynamics, fold-change of population in the first 14 days, and trace element detection. **a)** Schematic of the study design which *B. pseudomallei* strain 207a was inoculated into sterile water in 1994, showing nine initial concentrations, timeline and sampling strategies, including single-colony-picked and colony-sweep collections. **b)** Line charts demonstrated bacterial population dynamics of the 2023 replicated experiment (Log_10_CFU/mL) from inoculation to one year. **c)** Line charts depicted bacterial population dynamics of the 1994-original experiment (Log_10_CFU/mL), tracked 19-28 years post-inoculation. **d)** Bar charts showed fold-change in Log_10_CFU/mL between day 0 and day 14 across nine populations. **e)** The LC-MS/MS spectrum of sterile water used in the replication experiment with relative abundance over mass-to-charge ratio (m/z). Black peaks represent compounds detected in sterile water, red peaks indicate potential background contamination, grey highlights plastic residue (phthalic anhydride).

## Results

### Initial growth and long-term equilibrium under starvation

Since the experiment began in 1994, we have monitored both the viability and evolutionary trajectory of *B. pseudomallei* populations under nutrient-depleted conditions. Despite the wide range of initial inoculum densities (10^2^ to 10^10^ CFU/mL), viable cell counts across all tubes gradually converged to a median of ∼1 × 10^6^ CFU/mL by year 16 as previously reported^18^. From year 19 onward, this equilibrium has remained stable, with a median of ∼4.9 × 10^5^ CFU/mL (range: 0.79 x 10^5^–3.6 × 10^6^ CFU/mL), consistent with the establishment of a long-term steady-state under nutrient-depleted conditions (**Figure 1 b - c**). This steady-state population density is comparable to that reported for other Gram-negative bacteria, such as *Escherichia coli* and *Pseudomonas aeruginosa,* maintained in a similar *in vitro* system under prolonged starvation^28–30^. To characterise how populations initiated at different densities approach this equilibrium, we replicated the experiment in 2023 using the original experimental conditions. Population trajectories differed as a function of initial inoculum. High-density initial inocula (10^7^– 10^10^ CFU/mL) exhibited an initial increase, followed by a gradual decline in viable cell counts (**Figure 1b**), likely attributable to early use of carried-over nutrients and subsequent scavenging phase in which surviving cells exploit nutrients released from lysed neighbours^31,32^. Interestingly, two tubes inoculated at densities close to the eventual equilibrium (10^5^-10^6^ CFU/mL) showed a similar initial increase but stabilised at comparable population levels within 14 days of the inoculation. In contrast, tubes inoculated at much lower density (10^2^-10^4^ CFU/mL) ultimately reached similar equilibrium cell concentrations despite the absence of external nutrient input. Given the minimal original cell biomass, the observed 6.22-log_10_-fold increase in cell numbers (range: 6.18-6.25 log_10_-fold increase, **Figure 1d**) cannot be explained by dead-cell scavenging alone. Although reductive cell division accompanied by morphological changes from rod to cocci shape may contribute to population maintenance or expansion^18^, these observations raise the possibility that alternative nutrient sources were available within the experimental set up.

### Plastic-derived compounds as potential carbon sources under nutrient-limited conditions

We hypothesised that trace organic compounds — either introduced during water manipulation or incompletely removed by distillation — may have served as a carbon source to support early metabolism and growth. To test this, we performed liquid chromatography-tandem mass spectrometry (LC-MS/MS) on the sterile water used in the replicate experiment. Given the expected low abundance of trace compounds and potential contamination from the analytical instruments, we prepared serial dilutions of the sterile water using ultrapure Milli-Q water at four different concentrations. For each detected compound, we estimated the true intensity in sterile water relative to background contamination by fitting a multiple linear regression across dilution series. This approach allows us to quantify the relative abundance of trace compounds in the sample versus instrumental background (**Figure 1e**, **Supplementary Table 2**, **see Methods**). We detected 558 compounds in the sterile water with varying levels of potential contamination (**Supplementary Table 2**). Each compound was screened for known degradation pathways in *B. pseudomallei* or closely related *Burkholderia* species (**Supplementary method**).

Among these, phthalic anhydride (PA), a C_8_ phthalate-related compound, was detected at relatively high abundance in both the sterile water and on the instruments (**Figure 1e**). PA is a common environmental contaminant known to leach from plastic medical and laboratory materials^33^. *B. pseudomallei* and related *Burkholderia* species possess genes putatively homologous to those for phthalate importation and degradation pathways^34^, including *phtAabd phtBR* operon^35^ (**Supplementary figure 1**). These enzymes convert PA into protocatechuate and subsequently to β-ketoadipate^36^, which can be oxidised to acetyl-CoA which enters the tricarboxylic acid cycle, thereby linking phthalate catabolism to central cell metabolism. This metabolic pathway has also been described in other gram-negative bacteria, including *Acinetobacter* species^37^. Although PA was detected in both sterile water and the LC-MS/MS instrument background, its presence in the sterile water at the start of the experiment is consistent with a role in supporting initial growth of *B. pseudomallei*. Over time, additional PA — likely leached from the plastic tubes used in the original 1994 experiments (plastic was chosen over glassware in BSL3 for safety reasons) – could have served as a continued carbon source, sustaining the population during the long-term survival phase. While other compounds were also detected (**Figure 1e**), we cannot exclude the possibility that additional substrates contributed to survival under nutrient-depleted conditions. Nevertheless, no genomic or functional evidence currently supports the ability of *B. pseudomallei* to utilise these compounds as carbon sources.

### Minimal genomic changes detected in *B. pseudomallei* after prolonged nutrient limitation suggest dormancy and slow growth

We compared the genetic variations accumulated in descendant strains to the 1994 founder strain, which was sequenced using short-read and long-read technologies and assembled into complete chromosomes. Descendant strains were sequenced with short-read technology as single-colony representative isolates recovered from viability assays conducted in 2013 (n = 2), 2019 (n = 6), and 2022 (n = 170), covering all inoculation tubes (mean sequencing coverage 42x). To minimise sampling bias from single-colony picks, we also performed population-level short-read whole-genome sequencing on pooled plate-sweep colony mixtures from each 2022 tube (n = 9 bulk samples, mean coverage 87.1x). Using single-colony genomes alone, we detected one large 264.6-kb deletion, 55 small insertions or deletions (indels), and 171 single nucleotide polymorphisms (SNPs) across descendant genomes. Population-level sequencing recovered largely overlapping variants and detected additional variants not captured among the single-colony isolates, consistent with the presence of low-frequency or population-specific mutations (**Supplementary Figure 2**). When combining evidence from both approaches, we detected one large deletion, 73 indels and 249 SNPs, affecting 266,132 bp of the 7,123,344 bp genome (3.74%) and impacting 393 of 5,881 predicted protein-coding sequences (6.68%) (**Figure 2a**).

**Figure 2.**
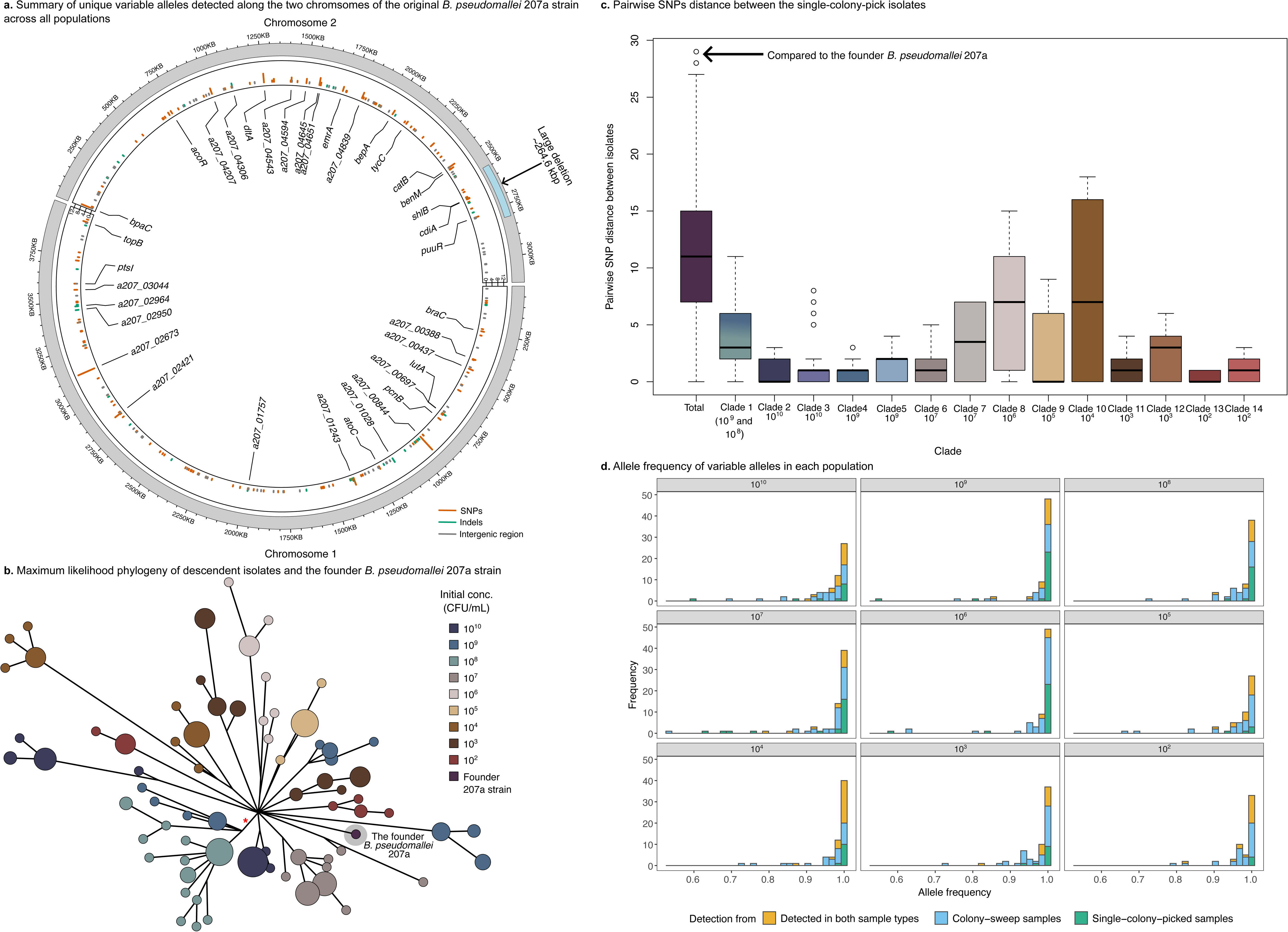
Distribution of genetic variants observed across nine populations **a)** Circular diagrams of unique genetic variants relative to the *B. pseudomallei* 207a founder isolate across two chromosomes, with bar charts summarising the distribution of variants counts across the populations. SNPs, indels, and intergenic variants are highlighted in orange, green, and grey, respectively. **b)** Maximum likelihood phylogeny of descendent isolates and the *B. pseudomallei* 207a founder isolate. The phylogenetic tree is illustrated in minimum spanning tree for better interpretation. Each tip is coloured by inoculum. A clade containing a homoplasic site which appears as a common branch is highlighted with a red star. **c)** Boxplots summarising pairwise SNP distances among isolates in this study. The distribution is depicted for the entire population and each clade. **d)** Bar charts of variant allele frequency observed in each population, with variants detected from single-colony-picked, colony-sweep and both methods, coloured green, blue, and yellow, respectively.

We next constructed a maximum likelihood phylogeny using whole-genome sequences of 170 single-colony isolates collected between 2013 and 2022, rooting the tree with the 1994 founder strain (**Figure 2b, Supplementary Figure 3**). The tree, supported by bootstrap values >70, resolved 14 genetically distinct clades, each corresponding to a specific inoculation tube, consistent with independent evolution under spatially isolated conditions. Pairwise SNP distances were low both between clades (median 11 bp) and within each clade (median 2 bp) (**Figure 2c**), limiting our ability to confidently assign variants detected by population-level sequencing to specific clades or incorporate bulk-sequenced samples directly into the phylogeny.

Using a recombination-removed phylogeny, we estimated tube-specific mutation rates (**Supplementary Figure 4**, **Supplementary method**). Incorporating known sampling dates, mutation rates were calculated based on the number of SNPs accumulated relative to the most recent common ancestor of the sampled isolates, yielding a mean mutational rate of 3.18 x 10^-8^ SNPs per site per year [95% CI: 2.90 x 10^-8^ – 3.46 x 10^-8^ SNPs per site per year]. Although we note the limited sample size, this rate is substantially lower than previous estimates derived from a chronic 16-year *B. pseudomallei* infection (1.7 x 10^-7^ SNPs per site per year^38^), supporting a model in which most cells likely entered an extremely slow-growing or dormant state, resulting in minimal replication and reduced mutation accumulation over nearly three decades.

### Parallel genetic adaptation revealed by dual approaches

We hypothesised that differences in initial inoculum density imposed distinct selective pressures, potentially leading to parallel but independent adaptive mutations in the population from low (10^2^-10^4^ CFU/mL), equilibrium stage (10^5^-10^6^ CFU/mL), and high (10^6^-10^10^ CFU/mL) initial inoculum. To test this, we applied two complementary approaches: (i) a phylogenetic-based method to identify homoplasic mutations arising independently in different clades from single-colony genomes, and (ii) a frequency-based method to detect recurrent mutations across pooled plate-sweep population sequencing, grouped by initial inoculum density (**Figure 2d & 3a, Supplementary Table 3**).

Using 200 high-confidence SNPs, we mapped variants onto a maximum-likelihood phylogeny to detect homoplasies (**Supplementary figure 3).** SNPs present in all descendant isolates were excluded, as they likely represent variants introduced during the founder strain passage prior to the initiation of the experiment or artefacts arising from founder sequencing (**Supplementary method**). This analysis identified three homoplasic SNPs affecting three genes, each occurring independently in multiple clades. In parallel, we performed frequency-based analysis of pooled population sequencing data to identify recurrent variants across experimental tubes. As with the phylogeny-based approach, SNPs and indels detected in all populations were excluded. In total, we identified an additional 79 SNPS and 21 indels co-detected in more than one tube from the pooled populations, with allele frequencies ranging from 53.3% to 100% per tube, collectively affecting 40 genes (**Supplementary Table 3**). Notably, 52.7% of recurrent variants were present at sub-fixation frequencies (<100%), indicating substantial within-population genetic heterogeneity (**Figure 2d**). A subset of these variants was associated with specific inoculum density ranges: low-density populations (10^2^-10^4^ CFU/mL) harboured 7 SNPs and 6 indels affecting 10 genes; high-density populations (10^7^-10^10^ CFU/mL) contained 15 SNPs and 8 indels affecting 17 genes; and equilibrium-density populations (10^5^ – 10^6^ CFU/mL) exhibited 10 SNPs and 3 indels affecting 12 genes. Overall, the frequency-based approach extended variants detection beyond single-colony genomes, increasing the number of detected variants from three SNPs in three genes to 249 SNPs and 73 indels across 194 genes Importantly, all three homoplasic SNPs identified using the phylogeny-based method were also detected by the frequency-based approach (**Supplementary Table 3**). These involved mutations in homologues of *qseC*, *ermA* and *rssB*. *qseC* encodes a quorum sensing sensor kinase. Phylogenetic-based method identified an L343R substitution occurring independently in high-density populations, while frequency-based approach detected the same variant and additionally identified a Q215_P220dup frameshift mutation in equilibrium-and high-density tubes, a D411G substitution in equilibrium-density tubes, and an L192P in low-density tubes. All variants are predicted to affect protein structure (**Supplementary Figure 5**). For *emrA*, encoding a component of a multidrug efflux pump, both phylogeny-and frequency-based approaches detected a E10D substitution, while the frequency-based analysis further identified a nonsynonymous T52P substitution in an equilibrium-density population (**Supplementary Figure 6**). Finally, *rssB*, a regulator of sigma factor proteolysis, exhibited an R142G nonsynonymous substitution within the PAS sensor domain in equilibrium-and high-density populations, which was independently detected by both approaches (**Supplementary Figure 7**). Together, these findings illustrate how complementary analytical strategies reveal recurrent, density-associated genetic changes, and provide convergent evidence for parallel adaptation under long-term nutrient-depleted conditions.

### Functions and examples of genes with the highest mutational burden under nutrient depletion

Among the 193 genes exhibiting recurrent mutations (**Figure 3a**), some recurrence could in principle arise by chance, particularly in longer genes that have higher probability of accumulating mutations. To distinguish non-random signals from gene-length effects, we applied a gene-length-corrected mutational burden test (**Figure 3b**). This analysis identified 27 genes with a significantly elevated mutational load (Poisson test, Bonferroni-adjusted *p-value* < 0.05, **Supplementary Table 4**). To independently validate these signals, we performed 10,000 permutation simulations in which a total number of observed mutations was preserved but randomly reassigned across all chromosomes. For each gene, the observed mutation count was compared with the null distribution generated by permutation, and a permutation p-value was calculated as the proportion of simulations in which the simulated mutation count equalled or exceeded the observed value. The same 27 genes identified by the mutational burden test were also significantly enriched in the permutation analysis (*p-value* < 0.05, **Supplementary Table 4**), providing independent support for non-random accumulation of mutations at these loci. We hereafter refer to these 27 genes as the high-burden set.

**Figure 3.**
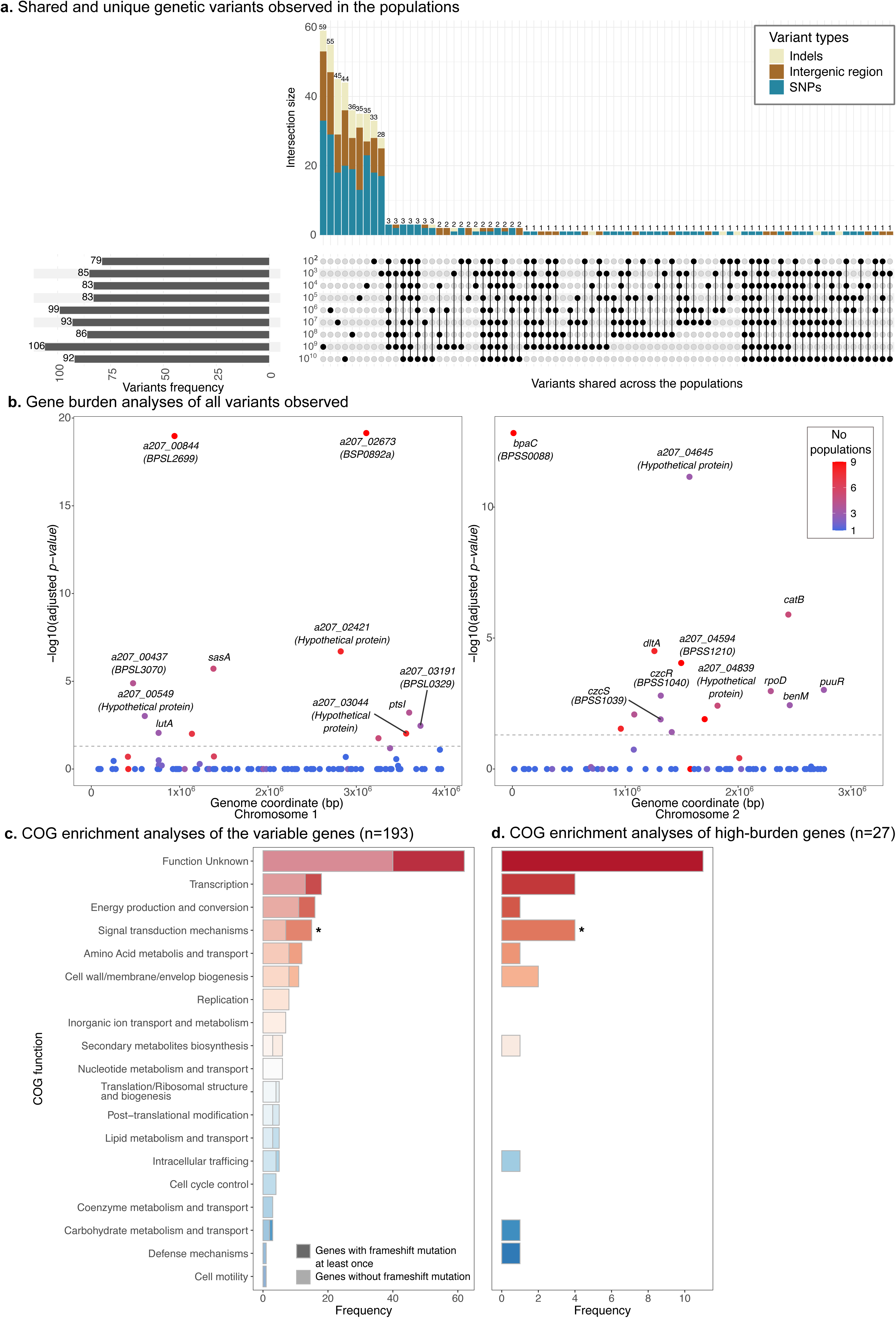
Parallel genetic adaptation after 28-years in sterile water **a)** Upset plot illustrating shared and unique genetic variants observed across nine inoculums. The types of genetic variants observed, SNP, Indel and intergenic variants, were highlighted in blue, cream, and brown. **b)** Manhattan plot across two chromosomes depicting genes with significantly elevated mutations compared to expected by chance under Poisson assumption across nine tubes. Points are coloured by the number of populations carrying the variants, from blue (unique to one population) to red (present in all nine). The dotted line denotes non-significant results after Bonferroni correction (adjusted *p-value* > 0.05) **c)** Stacked bar chart summarising the COG functional categories of genes with variable alleles. A proportion of genes in each COG function with frame-shift mutation at least once is highlighted. The star is denoted a significantly functional enrichment with *p-value* of < 0.05 by two-tailed fisher’s exact test. **d**)Bar chart summarising the COG functional categories of genes with significant mutational burdens. The star is denoted a significantly function enrichment with *p-value* of <0.05 by two-tailed fisher’s exact test.

We next assigned functional annotations to both the full set of 193 recurrently mutated genes and the subset of 27 high-burden genes (**Figure 3c-d**). Of the 193 genes, 131 could be classified into COG functional categories. Predominant functional classes (n > 5) included transcription, energy production and conversion, signal transduction, amino acid metabolism, cell envelope biogenesis, replication, ion transport, and secondary metabolite biosynthesis. Insertions or deletions predicted to cause frameshifts or premature stop codons - indicative of functional gene disruption – were widespread, occurring at least once in 53 of the 193 genes and spanning all functional categories. This pattern is consistent with the accumulation of loss-of-function mutations in genes that are dispensable under prolonged nutrient deprivation.

Signal transduction genes were significantly enriched among both the full set of 193 recurrently mutated genes and the 27 high-burden genes (two-tailed Fisher’s exact test, *p-value* = 2.61×10^-3^). Notably, signal transduction genes also exhibited a significantly higher frequency of disruptive mutations compared to other functional categories (two-tailed Fisher’s exact test, *p-value* = 0.0434, **Supplementary table 5**), suggesting preferential inactivation of signalling pathways that are no longer required during long-term nutrient limitation. For example, a homologue of the metal-sensing kinase *czcS* accumulated a frameshift mutation and a premature stop codon, both predicted to abolish protein function (**Supplementary Figure 8**). Several signal-transduction genes are represented by multiple paralogues in the *B. pseudomallei* genome, and these paralogues showed distinct mutational trajectories under extreme nutrient starvation. One paralogue of the quorum sensing kinase (*a207_02794*) carried a disruptive insertion together with two additional non-synonymous substitutions, whereas a second paralogue (*a207_00082*) accumulated a separate non-synonymous mutation (**Supplementary Figure 5**). Similarly, the sensory kinase *sasA* exhibited differential mutational patterns across three paralogues: one paralogue (*a207_01241*) accumulated one insertion and five non-synonymous substitutions, a second paralogue (*a207_04759*) carried a frameshift mutation and additional non-synonymous changes, while a third paralogue (*a207_04904*) accumulated only synonymous substitutions (**Supplementary Figure 9**). Notably, these mutations were differentially distributed across high-and low-density populations (**Supplementary Table 3**), consistent with population-density–dependent modulation of signal transduction.

In contrast, envelope-associated proteins and adhesins tended to accumulate predominantly non-synonymous mutations rather than disruptive variants. Although not statistically enriched, these patterns suggest functional remodelling rather than gene inactivation. A homologue of *BPSL0892a*, encoding a putative lipoprotein, accumulated 14 mutations, including five non-synonymous substitutions, all located within annotated protein domains (**Supplementary Figure 10**). Likewise, a homologue of *BPSS0088*, a trimeric autotransporter, accumulated 18 mutations, including nine non-synonymous changes within functional domains (**Supplementary Figure 11**), potentially reflecting altered cell–cell or surface interactions during prolonged starvation. Together, these results indicate that under long-term nutrient depletion, *B. pseudomallei* repeatedly targets signal transduction pathways for disruption, while selectively modifying surface-associated and biosynthetic functions, consistent with a transition toward a low-activity, survival-oriented physiological state.

## Conclusion and discussion

Our study provides a unique resource and framework for understanding the adaptive strategies that enable long-term survival of environmental bacteria under extreme nutrient depletion, using *B. pseudomallei* as a model. We report on *B. pseudomallei* maintained in sterile water for nearly three decades and used single-colony-picks and colony-sweep approaches in tandem to provide a more comprehensive picture of the adaptation process occurring under extreme nutrient limitation. Our results show that bacterial adaptation under prolonged nutrient deprivation relies on a combination of early-stage metabolic flexibility and long-term genetic adaptation, with many mutations fixed independently across subpopulations, reflecting parallel evolution.

Genetic characterisation highlights classes of mutations ranging from nonsynonymous to indels or frameshift mutations, which are likely loss-of-function. These mutations were a dominant feature of long-term adaptation. Under nutrient-limited conditions, selection favours the inactivation of non-essential or metabolically costly genes – such as those involved in signal transduction mechanisms – as an energy conservation strategy. These mutations reduce cellular expenditure and enhance survival at the population level. Notably, loss-of-function events were tube-or clade-restricted, contributing to phenotypic heterogeneity, where subpopulations may adopt diverse states (e.g. persistence or dormancy) to withstand unpredicted environmental stress. This pattern mirrors observations in chronic infections, such as *Pseudomonas aeruginosa* in cystic fibrosis^40^, where survival – rather than growth – is selected.

Metabolic flexibility was critical during early adaptation. *B. pseudomallei* can utilise alternative carbon sources when common nutrients are scarce^21^, including trace anthropogenic compounds such as plastic-derived residues. This likely facilitated early growth in low-inoculum tubes and contributed to sustained biomass under nutrient deprivation. From a public health perspective, this capacity may enhance bacterial persistence in non-natural environments, such as plumbing systems^41^, which can harbour synthetic chemicals^42^ and organic pollutants^43^. Conversely, in synthetic biology, this metabolic versatility could potentially be harnessed for bioremediation or plastic waste management.

While our study is among the most comprehensive genomic analyses of long-term bacterial survival under starvation to date, several limitations exist. The original 1994 experiment had irregular sampling intervals, incomplete environmental records (e.g. fluctuating lab temperatures), some loss of experimental data, and the limited biomass remaining in each tube restricted full biochemical characterisation without compromising the original setup. A replication of the experiment in 2022 helped to supplement these gaps, although exact replication of original conditions was not possible. Nonetheless, combining new and archived data from 1994, 2013, 2019, and 2022 allowed us to capture both early survival events and long-term evolutionary dynamics.

Overall, our findings demonstrate that *B. pseudomallei* combines metabolic flexibility and long-term genetic adaptation to persist for decades in nutrient-poor environments. This highlights the bacterium’s remarkable ecological resilience and provides insights into the energy-saving survival strategies that sustain its persistence in the environment and potentially during host infection.

## Method and materials

### Study design

In 1986, *Burkholderia pseudomallei* strain 207a was recovered from a blood sample of a rice farmer admitted to Sunpasitthiprasong Hospital, Ubon Ratchathani Province, Northeast Thailand and subsequently preserved at-80°C in trypticase soy broth (TSB) containing 15% glycerol. In 1994, Wuthiekanun *et al*, sub-cultured this *B. pseudomallei* strain onto enrichment Columbia agar and inoculated it into 9 mL sterile triple-filtered distilled water to obtain a master concentration of 1×10^10^ CFU/mL in a plain plastic tube^16^. They then performed a serial dilution resulting in nine tubes with initial concentrations of 10^2^ - 10^10^ CFU/mL. The tubes were tightened and then loosened by a half turn to allow the bacteria to perform minimal aerobic respiration. These nine tubes were maintained in a lab cabinet at ambient temperature in a secure biosafety level-3 laboratory. In recent years, an aliquot was removed from each tube and cultured to check for bacterial viability. We collected 170 isolates from these plates using a single-colony-pick in 2013 (n=2), 2019 (n=6), and 2022 (n=161) to further investigate bacterial adaptation through genetic studies. To comprehensively capture the genetic diversity within the populations, we further collected 9 pooled populations from each tube using a colony-sweep method. These isolates were preserved at-80°C in trypticase soy broth (TSB) containing 15% glycerol before DNA extraction in the year 2022.

The growth rate of strain 207a following exposure to distilled water would demonstrate early adaptation to nutrient deprivation. However, the original 1994 records covering this early adaptation are missing. Therefore, we replicated the experiment under similar conditions. Briefly, the founder 207a strain was inoculated into selective Ashdown media before re-subculturing onto enrichment Columbia media plate. The colonies of the original 207a isolates were collected directly from the Columbia agar plate into 30 mL sterile deionised (DI) water and adjusted into 10^10^ CFU/mL before performing serial dilution into nine concentrations (between 10^2^ to 10^10^ CFU/mL). The tubes were kept in the lab cabinet at ambient temperature (**Figure 1a**). We then observed each tube for changes in bacterial cell counts on day 1, day 3, day 7, and weekly thereafter. Three replicates have been performed. Furthermore, to enhance the ability to trace the genetic and evolutionary change, a sample will be collected every three months for future studies.

### Trace element analyses of sterile water

Given the bacterial population expansion observed in the first 14 days upon inoculation into the nutrient-free environment, we further determined the possible nutrient sources in deionised water used in this replicated experiment. We identified the trace elements in DI water using an untargeted metabolomics approach via Liquid Chromatography - Tandem Mass Spectrometry (LC-MS/MS). DI water and LC-MS water were compared to assess trace element contamination, with LC-MS water serving as the control. For each water type, 500 mL was freeze-dried and reconstituted in the mobile phase. To confirm the origin of detected compounds, the reconstituted sample was diluted in mobile phase at volume ratios of 1:1, 1:2, 1:4, and 1:9 (sample:mobile phase, v/v). These dilutions were prepared not for quantification, but to verify whether the intensity of true water-derived compounds decreased proportionally with dilution—supporting their presence in the original water. Each sample both undiluted and diluted was injected into the LC-MS/MS system four times to ensure technical reproducibility.

Untargeted metabolomics profiling was performed to identify trace chemical constituents in sterile water and distinguish authentic sample-derived features from instrument-associated background signals. Samples were analysed using reverse-phase liquid chromatography coupled to a high-resolution QTOF 5600+ mass spectrometer (Sciex). A 5 µL injection volume was used and chromatographic separation was achieved on a UPLC Acquity HSS T3 C18 column (1.8 μm, 2.1×100 mm) with a matched trap column, maintained at 40 °C. Mobile phases consisted of water with 0.1% formic acid (A) and acetonitrile with 0.1% formic acid (B), delivered at 0.3 mL min⁻¹ under the following linear gradient program: 0-1.5 min, 1%B; 1.5-3.5 min, 1→15%B; 3.5-6.5 min, 15→50%B; 6.5-9.5 min, 50→95%B; 9.5-12.0 min, 95%B; 12.0-12.1 min, 95→1% B; and 12.1-16.0 min, 1%B. Data acquisition was performed using Analyst Software v1.7 (SCIEX), with spectra collected in both positive and negative electrospray ionisation modes. Instrument settings for the TripleTOF 5600 were as follows: IonSpray voltage of +5500 V in positive mode and-4500 V in negative mode; curtain gas set to 30 psi for +ESI and 25 psi for-ESI; ion source gas 1 (GS1) of 45 psi, ion source gas 2 (GS2) of 40 psi; source temperature 450 °C; and a collision energy ramp of 15-45 V. Data were acquired in the information-dependent acquisition (IDA) mode, consisting of a TOF-MS survey scan followed by ten dependent product-ion scans in high-sensitivity mode with dynamic background subtraction enabled. TOF-MS scans were collected over m/z 100-1000, and MS/MS product-ion scans over m/z 50-1000.

The raw LC-MS/MS spectra were first centroided and converted into mzML format using ProteoWizard’s MSConvertGUI^44^. Metabolic features were then captured from the pre-processed spectra by integrating MS1 (TOF-MS scan) and MS2 (MS/MS product-ion scan) spectral data using an R package MetaboAnalystR V4.0^45^. Compound identification was done by which matching processed MS2 data against reference spectra databases including; HMDB^46^, MoNA Series^47^, LipidBlast^48^, MassBank^49^, GNPS^50^, LipidBank^51^, MINEs^52^, LipidMAPs^53^, and KEGG^54^. Identification results were reported with matching scores considering m/z spectra, retention time, isotope and MS2 similarity. The similarity scores range between 0 (no match) to 100 (perfect match). To control for potential misidentification of the metabolites observed, we applied a 70% similarity score cut-off to remove any artefacts or false positives. This cut-off is chosen following common practice in a spectra-library match metabolomic workflow^55^.

### Quantitative comparison of compound intensities in sterile water versus instrument-derived contaminants

To distinguish authentic chemical signatures, present in sterile water from background signals originating from the analytical system, ultrapure MiliQ water was analysed as a procedural control. Compounds detected in MiliQ water were therefore interpreted as instrument-or solvent-derived contaminants. We applied a linear mixed-effects model across four dilution levels to deconvolute true sample-derived signals from background noise, expressed as:

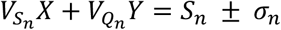

*X* represents the estimated abundance of compounds intrinsic to the sterile water, while *Y* represents the abundance attributable to instrument-derived contamination as measured in MiliQ water. The coefficients *V(Sₙ)* and *V(Qₙ)* correspond to the applied dilution factors for each sample type. The term *Sₙ* captures the raw LC-MS/MS signal intensity, with *σₙ* reflecting the associated measurement uncertainty. This modelling framework enables robust separation of genuine water-borne compounds from artefactual background features introduced during sample handling or ionisation.

### Bacterial growth, DNA extraction, sequencing, and quality control

The isolates from the original experiment were cultured on the selective Ashdown media, followed by subculture on an enrichment Columbia media in a biosafety level 3 laboratory. A 1 ml loopful of *B. pseudomallei* colonies was collected before being washed in 1 mL of sterile PBS solution. The bacterial mixture was spun down at 21,200 rcf for 5 mins before the supernatant was removed. DNA extraction was performed using QIAamp DNA mini kit (Qiagen, Germany). The purity of extracted DNA was assessed by A260/280 ratio using Nanodrop (Thermo Scientific, U.S.A). Whole-genome sequencing was performed using Illumina Novaseq with 150 bp read length under a collaboration with Wellcome Sanger Institute, UK.

### Colony-sweep sample collection and data

To identify genetic variations that we might have missed from the single colony picks^14–16^, we employed a colony-sweep technique for all 9 inoculum concentrations (n=9) followed by DNA extraction using the column-based technique as described above. We then sequenced the pooled DNA using short read whole genome sequencing. The library preparation and sequencing were performed using Illumina Novaseq with 150 bp paired end runs at high depth resulting in 2,653,331 (range 2,419,700 - 2,984,290) reads per sample (**Supplementary table 1**).

We further performed taxonomic classification to identify any potential contamination in each sample, both for isolates obtained from single-colony picks and pooled DNA from mixed plate-sweep colonies using Kraken V1.1.1^56^, CheckM2 V1.0.1^57^ and FastANI V1.3^58^ to ensure accurate species-level identification, genome completeness, and to detect any cross-species contamination or mis-assemblies.

### Genome assembly and annotation

The genome of the original 207a strain was sequenced using both short-read Illumina Novaseq and long-read Oxford Nanopore Technology (ONT). The library preparation step for the ONT was performed as per the manufacture’s instruction using Ligation sequencing kit. An additional filter step was performed on long-read sequences to remove low quality reads by Filtlong V0.2.1 with options --min_length 1000 --keep_percent 95^59^. We then assembled the genome of the original 207a strain using Unicycler V0.5.0^60^ and used it as the reference strain for variant calling. The quality of the assembled sequences was inspected by Quast V5.0.2^61^. Gene predictions and annotations of assemblies were performed using Prokka V1.14.5^62^. Given many genes were annotated as hypothetical proteins, COG and GO-term analyses were performed to further categorise the function of annotated proteins by eggNOG-mapper V2.1.11^63^ and InterPro V100.0^64^. A hybrid-assembly of the original reference 207a strain yielded two complete chromosomes with a predicted 5,787 coding sequences (CDS), and a total length of the whole genome of 7,127,344 bps (Chromosome 1 - 3,997,571 bps and Chromosome 2 - 3,129,773 bps). The average GC content of the 207a strain is 68.27 which is similar to the reference genome K96243 strain from Northeast Thailand (**Supplementary figure 12**).

*De novo* assembly of short-reads of 207a descendant isolates was performed using either Velvet V1.2.10^65^ for the samples collected in 2013 and 2019, while the samples collected in 2022 were assembled with SPADES V3.15.5^66^. The differences in assembly used were due to the upgraded version of the assembly pipeline. To assess the quality of our assemblies, we further employed a quality control strategy as described above. *De novo* assembly of the descendent isolates led to genomes with a median of 74 contigs (range of 61-163) and an average total length of 7,062,204 bps (min = 6,674,100 bps and max = 7,084,479 bps) (**Supplementary Table 1**).

### Sequencing mapping and pseudo-alignment generation

Short reads of 207a descendant isolates were mapped against the parental 207a strain and a pseudo-alignment was constructed using a custom mapping pipeline (multiple_mappings_to_bam-1.6.py^67^) with bwa-MEM (v.0.7.17)^68^ as the aligner. A visual inspection of the alignment was performed using Seaview^69^ to check for unbiased mapping.

### Population structure estimated by phylogenetic tree building and ancestral reconstruction

Recombination was detected and removed from the whole-genome pseudo-alignment using Gubbins V3.2.1^70^. SNPs between the isolates in the alignment were identified using SNP-sites V2.5.1^71^ and used to construct a maximum likelihood phylogeny with 1000 ultra-bootstraps support using IQ-TREE V1.6.12^72^ with ModelFinder option. The best-fit model was selected as K3P+ASC with each branch possessing at least 70% bootstrap support. The phylogenetic tree and alignment were further analysed for homoplasy using HomoplasyFinder R package^73^ to identify mutations which have evolved independently in separate evolutionary lineages due to convergent evolution. The final phylogenetic tree is illustrated as a minimum spanning tree produced by GrapeTree V1.5^74^. To explore the order of mutations required for adaptation in sterile water, SNPs were reconstructed onto each branch of the phylogeny using Treetime V0.11.3^75^ with ancestral option. This ancestral option was chosen to infer the maximum likelihood of the ancestral nucleotide sequences along the phylogeny utilising a General Time Reversible model.

### Genetic variants detection of single-colony-picked and colony-sweep samples

Genetic variations from single-colony-picked isolates, such as insertions and deletions (indels), were further investigated using Snippy V4.626^76^ with the previously hybrid-assembled 207a strain as a reference. Given all isolates are descendants of 207a, the counts of variations observed in each population were then corrected based on the known phylogenetic relationships. Briefly, we identified the members of each internal node in the previously constructed phylogeny. The genetic variants observed in all samples from each internal node are counted only once, reducing bias in the variants observed across the populations.

To capture minority variants present in the colony-sweep samples, we applied our custom analysis pipeline and stringent filtering criteria. Short reads from the colony-sweep samples were mapped to the parental 207 strain reference genome using BWA V7.1^68^ and BCFtools V1.9^77^ for variant detection. Filtering criteria were applied to minimise false positives due to sequencing errors as previously described by Bryant et al^78^. Briefly, the variants had to be supported by at least 5 reads, with a minimum of 2 reads on each strand, and a strand bias P-value threshold of 0.05. The allele frequency calculated from BCFtools was used to infer the subclonal populations within the colony-sweep populations. The variant effect was annotated using SNPEff^79^.

### Mutational burden analysis and functional identification

Taking together the variants detected from single-colony-picked and colony-sweep isolates, we further assessed the mutational burden of every gene using two independent approaches: a Poisson test with adjusted *p-value* < 0.05 and permutation test with 10,000 stimulations when mutations randomly distributed across the two chromosomes. The permutation test was performed by calculating the proportion of simulations in which the simulated mutation counts were equal to or greater than the observed count with significance assessed at < 0.05. Further, to elucidate the protein function of genes with adaptive signatures, we employed a protein structure prediction method, AlphaFold3^80^, using a default setting and random seed to identify protein structural adaptation. The protein structures were visualised and evaluated using Mol*Viewer^81^. In addition to the structural changes, we utilised InterProScan^82^ to predict the functional domains where the genetic variants localised to infer their functional implications for the proteins.

### Data visualisation and statistical analyses

Visualisation and statistical analyses were performed in R (version 4.5.0). Where appropriate, we used the Bonferroni method to adjust the *p-value* for multiple comparisons.

## Data availability

The genome sequence data generated in this study are publicly available in the European Nucleotide Archive (ENA) under the study numbers PRJEB35787, PRJEB48720 and PRJEB107654. Accession numbers for individual genomes are provided in **Supplementary table 1.** All source data is publicly available at https://doi.org/10.6084/m9.figshare.31236106.

## Acknowledgements

This paper is dedicated to the memory of Professor Nicholas J White. CCho was funded by the Prince Mahidol PhD Studentship. CChe was funded by the Wellcome International Intermediate Fellowship (216457/Z/19/Z), the Sanger International Fellowship, and the University of Oxford Nuffield Department of Medicine Career Development Scheme. This research was funded in part by the Wellcome Trust (220211 and 206194). We acknowledge the use of AlphaFold Server (DeepMind) for structure prediction. Ongoing use of the output is subject to the AlphaFold Server Terms of Use. For the purpose of Open Access, the author has applied a CC BY public copyright license to any Author Accepted Manuscript version arising from this submission. The funders had no role in study design, data collection and analysis, decision to publish, or preparation of the manuscript.

## Author contribution

CChe, VW, and JP contributed equally. NJW, VW, and CChe conceived the study. CChe, VW, PT, JTa designed the trace element detection of sterile water study. CChe, NJW, VW, JP, CCho, DL, JC, NPJD and NRT designed the study. CChe secured funding. PM and SL collected, identified bacterial isolates, and curated associated data. KS, SA, and PT processed the water samples. KC did the LC-MS/MS analyses. CChe, JP, NRT performed short-read sequencing while JTh and EB performed long-read sequencing. NRT and JP contributed software tools. CCho performed bioinformatics analyses. VRB and JAL performed protein structure prediction. CCho, CChe, and JP interpret the analyses. All authors had full access to all data in the study and had final responsibility for the decision to submit for publication. CCho wrote the first draft. CCho, CChe and JP reviewed, edited, and wrote revision drafts. All authors read and approved the manuscript.

## References

1. Currie, B. J. et al. The Darwin Prospective Melioidosis Study: a 30-year prospective, observational investigation. Lancet Infect. Dis. 21, 1737–1746 (2021).

2. Chantratita, N. et al. Characteristics and one year outcomes of melioidosis patients in Northeastern Thailand: a prospective, multicenter cohort study. Lancet Reg. Health - Southeast Asia 9, (2023).

3. Mukhopadhyay, C., Shaw, T., Varghese, G. M. & Dance, D. A. B. Melioidosis in South Asia (India, Nepal, Pakistan, Bhutan and Afghanistan). Trop. Med. Infect. Dis. 3, 51 (2018).

4. Lichtenegger, S. et al. Melioidosis in Mali: a retrospective observational study. Lancet Glob. Health 13, e1964–e1972 (2025).

5. Petras, J. K. et al. Locally Acquired Melioidosis Linked to Environment — Mississippi, 2020–2023. N. Engl. J. Med. 389, 2355–2362 (2023).

6. Kingsley, P. V., Leader, M., Nagodawithana, N. S., Tipre, M. & Sathiakumar, N. Melioidosis in Malaysia: A Review of Case Reports. PLoS Negl. Trop. Dis. 10, e0005182 (2016).

7. Wiersinga, W. J. et al. Melioidosis. Nat. Rev. Dis. Primer 4, 17107 (2018).

8. Clayton, A. J., Lisella, R. S. & Martin, D. G. Melioidosis: a serological survey in military personnel. Mil. Med. 138, 24–26 (1973).

9. Larsen, E., Smith, J. J., Norton, R. & Corkeron, M. Survival, Sublethal Injury, and Recovery of Environmental Burkholderia pseudomallei in Soil Subjected to Desiccation. Appl. Environ. Microbiol. 79, 2424–2427 (2013).

10. Inglis, T. J. J. & Sagripanti, J.-L. Environmental Factors That Affect the Survival and Persistence of Burkholderia pseudomallei. Appl. Environ. Microbiol. 72, 6865–6875 (2006).

11. Chen, Y.-L., et al. The concentrations of ambient Burkholderia pseudomallei during typhoon season in endemic area of melioidosis in Taiwan. PLoS Negl. Trop. Dis. 8, e2877 (2014).

12. Yip, T.-W. et al. Endemic Melioidosis in Residents of Desert Region after Atypically Intense Rainfall in Central Australia, 2011. Emerg. Infect. Dis. 21, 1038–1040 (2015).

13. Chen, P.-S., et al. Airborne Transmission of Melioidosis to Humans from Environmental Aerosols Contaminated with B. pseudomallei. PLoS Negl. Trop. Dis. 9, e0003834 (2015).

14. Chapple, S. N. J. et al. Whole-genome sequencing of a quarter-century melioidosis outbreak in temperate Australia uncovers a region of low-prevalence endemicity. *Microb*. Genomics 2, e000067 (2016).

15. Inglis, T. J. et al. Interaction between Burkholderia pseudomallei and Acanthamoeba species results in coiling phagocytosis, endamebic bacterial survival, and escape. Infect. Immun. 68, 1681–1686 (2000).

16. Wuthiekanun, V., Smith, M. D. & White, N. J. Survival of Burkholderia pseudomallei in the absence of nutrients. Trans. R. Soc. Trop. Med. Hyg. 89, 491 (1995).

17. Palasatien, S., Lertsirivorakul, R., Royros, P., Wongratanacheewin, S. & Sermswan, R. W. Soil physicochemical properties related to the presence of *Burkholderia pseudomallei*. Trans. R. Soc. Trop. Med. Hyg. 102, S5–S9 (2008).

18. Pumpuang, A. et al. Survival of Burkholderia pseudomallei in distilled water for 16 years. Trans. R. Soc. Trop. Med. Hyg. 105, 598–600 (2011).

19. Limmathurotsakul, D. et al. Predicted global distribution of Burkholderia pseudomallei and burden of melioidosis. Nat. Microbiol. 1, 15008 (2016).

20. Hantrakun, V. et al. Soil Nutrient Depletion Is Associated with the Presence of Burkholderia pseudomallei. Appl. Environ. Microbiol. 82, 7086–7092 (2016).

21. Chewapreecha, C. et al. Co-evolutionary Signals Identify *Burkholderia pseudomallei* Survival Strategies in a Hostile Environment. Mol. Biol. Evol. 39, msab306 (2022).

22. Seng, R. et al. Genetic diversity, determinants, and dissemination of Burkholderia pseudomallei lineages implicated in melioidosis in Northeast Thailand. Nat. Commun. 15, 5699 (2024).

23. Holden, M. T. G. et al. Genomic plasticity of the causative agent of melioidosis, Burkholderia pseudomallei. Proc. Natl. Acad. Sci. 101, 14240–14245 (2004).

24. Spring-Pearson, S. M. et al. Pangenome Analysis of Burkholderia pseudomallei: Genome Evolution Preserves Gene Order despite High Recombination Rates. PLoS ONE 10, e0140274 (2015).

25. Nandi, T., et al. *Burkholderia pseudomallei* sequencing identifies genomic clades with distinct recombination, accessory, and epigenetic profiles. Genome Res. 25, 129–141 (2015).

26. Chewapreecha, C. et al. Global and regional dissemination and evolution of Burkholderia pseudomallei. Nat. Microbiol. 2, 16263 (2017).

27. Shaw, T. et al. Environmental Factors Associated With Soil Prevalence of the Melioidosis Pathogen Burkholderia pseudomallei: A Longitudinal Seasonal Study From South West India. Front. Microbiol. 13, 902996 (2022).

28. Machreki, Y., Kouidhi, B., Machreki, S., Chaieb, K. & Sáenz, Y. Analysis of a long term starved *Pseudomonas aeruginosa* ATCC27853 in seawater microcosms. Microb. Pathog. 134, 103595 (2019).

29. Himeoka, Y., Gummesson, B., Sørensen, M. A., Svenningsen, S. L. & Mitarai, N. Distinct Survival, Growth Lag, and rRNA Degradation Kinetics during Long-Term Starvation for Carbon or Phosphate. mSphere 7, e01006–21 (2022).

30. Zion, S., Katz, S. & Hershberg, R. Escherichia coli adaptation under prolonged resource exhaustion is characterized by extreme parallelism and frequent historical contingency. PLOS Genet. 20, e1011333 (2024).

31. Rozen, D. E., Philippe, N., Arjan de Visser, J., Lenski, R. E. & Schneider, D. Death and cannibalism in a seasonal environment facilitate bacterial coexistence. Ecol. Lett. 12, 34–44 (2009).

32. Gray, D. A. et al. Extreme slow growth as alternative strategy to survive deep starvation in bacteria. Nat. Commun. 10, 890 (2019).

33. Wang, W. & Kannan, K. Leaching of Phthalates from Medical Supplies and Their Implications for Exposure. Environ. Sci. Technol. 57, 7675–7683 (2023).

34. Morya, R., Salvachúa, D. & Thakur, I. S. *Burkholderia*: An Untapped but Promising Bacterial Genus for the Conversion of Aromatic Compounds. Trends Biotechnol. 38, 963–975 (2020).

35. Stingley, R. L., Brezna, B., Khan, A. A. & Cerniglia, C. E. Novel organization of genes in a phthalate degradation operon of Mycobacterium vanbaalenii PYR-1. Microbiology 150, 3749–3761 (2004).

36. Li, J., Zhang, J., Yadav, M. P. & Li, X. Biodegradability and biodegradation pathway of di-(2-ethylhexyl) phthalate by *Burkholderia pyrrocinia* B1213. Chemosphere 225, 443–450 (2019).

37. Xu, J., Lu, Q., de Toledo, R. A. & Shim, H. Degradation of di-2-ethylhexyl phthalate (DEHP) by an indigenous isolate *Acinetobacter* sp. SN13. Int. Biodeterior. Biodegrad. 117, 205–214 (2017).

38. Pearson, T. et al. Pathogen to commensal? Longitudinal within-host population dynamics, evolution, and adaptation during a chronic >16-year Burkholderia pseudomallei infection. PLOS Pathog. 16, e1008298 (2020).

39. Pakdeerat, S. et al. Benchmarking CRISPR-BP34 for point-of-care melioidosis detection in low-income and middle-income countries: a molecular diagnostics study. Lancet Microbe 5, e379–e389 (2024).

40. Weimann, A. et al. Evolution and host-specific adaptation of Pseudomonas aeruginosa. Science 385, eadi0908 (2024).

41. Pakdeerat, S. et al. CRISPR-based environmental detection of Burkholderia pseudomallei identifies sanitation gaps and melioidosis risk in northeast Thailand. 2024.11.21.24317607 Preprint at 10.1101/2024.11.21.24317607 (2025).

42. Fang, W., Peng, Y., Muir, D., Lin, J. & Zhang, X. A critical review of synthetic chemicals in surface waters of the US, the EU and China. Environ. Int. 131, 104994 (2019).

43. Chen, M., Niu, Z., Zhang, X. & Zhang, Y. Pollution characteristics and health risk of sixty-five organics in one drinking water system: PAEs should be prioritized for control. Chemosphere 350, 141171 (2024).

44. Chambers, M. C. et al. A cross-platform toolkit for mass spectrometry and proteomics. Nat. Biotechnol. 30, 918–920 (2012).

45. Pang, Z. et al. MetaboAnalystR 4.0: a unified LC-MS workflow for global metabolomics. Nat. Commun. 15, 3675 (2024).

46. Wishart, D. S. et al. HMDB 5.0: the Human Metabolome Database for 2022. Nucleic Acids Res. 50, D622–D631 (2022).

47. Hilbig, M. & Rarey, M. MONA 2: A Light Cheminformatics Platform for Interactive Compound Library Processing. J. Chem. Inf. Model. 55, 2071–2078 (2015).

48. Kind, T. et al. LipidBlast in silico tandem mass spectrometry database for lipid identification. Nat. Methods 10, 755–758 (2013).

49. Horai, H. et al. MassBank: a public repository for sharing mass spectral data for life sciences. J. Mass Spectrom. JMS 45, 703–714 (2010).

50. Wang, M. et al. Sharing and community curation of mass spectrometry data with Global Natural Products Social Molecular Networking. Nat. Biotechnol. 34, 828–837 (2016).

51. Watanabe, K., Yasugi, E. & Oshima, M. How to Search the Glycolipid data in “LIPIDBANK for Web”, the Newly Developed Lipid Database in Japan. Trends Glycosci. Glycotechnol. 12, 175–184 (2000).

52. Jeffryes, J. G. et al. MINEs: open access databases of computationally predicted enzyme promiscuity products for untargeted metabolomics. J. Cheminformatics 7, 44 (2015).

53. Conroy, M. J. et al. LIPID MAPS: update to databases and tools for the lipidomics community. Nucleic Acids Res. 52, D1677–D1682 (2024).

54. Kanehisa, M., Furumichi, M., Sato, Y., Matsuura, Y. & Ishiguro-Watanabe, M. KEGG: biological systems database as a model of the real world. Nucleic Acids Res. 53, D672–D677 (2025).

55. Rainer, J. et al. A Modular and Expandable Ecosystem for Metabolomics Data Annotation in R. Metabolites 12, 173 (2022).

56. Wood, D. E. & Salzberg, S. L. Kraken: ultrafast metagenomic sequence classification using exact alignments. Genome Biol. 15, R46 (2014).

57. Chklovski, A., Parks, D. H., Woodcroft, B. J. & Tyson, G. W. CheckM2: a rapid, scalable and accurate tool for assessing microbial genome quality using machine learning. Nat. Methods 20, 1203–1212 (2023).

58. Jain, C., Rodriguez-R, L. M., Phillippy, A. M., Konstantinidis, K. T. & Aluru, S. High throughput ANI analysis of 90K prokaryotic genomes reveals clear species boundaries. Nat. Commun. 9, 5114 (2018).

59. GitHub - rrwick/Filtlong: quality filtering tool for long reads. https://github.com/rrwick/Filtlong.

60. Wick, R. R., Judd, L. M., Gorrie, C. L. & Holt, K. E. Unicycler: Resolving bacterial genome assemblies from short and long sequencing reads. PLOS Comput. Biol. 13, e1005595 (2017).

61. Gurevich, A., Saveliev, V., Vyahhi, N. & Tesler, G. QUAST: quality assessment tool for genome assemblies. Bioinformatics 29, 1072–1075 (2013).

62. Seemann, T. Prokka: rapid prokaryotic genome annotation. Bioinforma. Oxf. Engl. 30, 2068–2069 (2014).

63. Cantalapiedra, C. P., Hernández-Plaza, A., Letunic, I., Bork, P. & Huerta-Cepas, J. eggNOG-mapper v2: Functional Annotation, Orthology Assignments, and Domain Prediction at the Metagenomic Scale. Mol. Biol. Evol. 38, 5825–5829 (2021).

64. Paysan-Lafosse, T. et al. InterPro in 2022. Nucleic Acids Res. 51, D418–D427 (2023).

65. Zerbino, D. R. & Birney, E. Velvet: Algorithms for de novo short read assembly using de Bruijn graphs. Genome Res. 18, 821–829 (2008).

66. Bankevich, A. et al. SPAdes: A New Genome Assembly Algorithm and Its Applications to Single-Cell Sequencing. J. Comput. Biol. 19, 455–477 (2012).

67. GitHub - sanger-pathogens/bact-gen-scripts. https://github.com/sanger-pathogens/bact-gen-scripts.

68. Li, H. Aligning sequence reads, clone sequences and assembly contigs with BWA-MEM. Preprint at 10.48550/ARXIV.1303.3997 (2013).

69. Gouy, M., Guindon, S. & Gascuel, O. SeaView Version 4: A Multiplatform Graphical User Interface for Sequence Alignment and Phylogenetic Tree Building. Mol. Biol. Evol. 27, 221–224 (2010).

70. Croucher, N. J. et al. Rapid phylogenetic analysis of large samples of recombinant bacterial whole genome sequences using Gubbins. Nucleic Acids Res. 43, e15 (2015).

71. Page, A. J. et al. SNP-sites: rapid efficient extraction of SNPs from multi-FASTA alignments. *Microb*. Genomics 2, e000056 (2016).

72. Minh, B. Q. et al. IQ-TREE 2: New Models and Efficient Methods for Phylogenetic Inference in the Genomic Era. Mol. Biol. Evol. 37, 1530–1534 (2020).

73. Crispell, J., Balaz, D. & Gordon, S. V. HomoplasyFinder: a simple tool to identify homoplasies on a phylogeny. *Microb*. Genomics 5, e000245 (2019).

74. Zhou, Z. et al. GrapeTree: visualization of core genomic relationships among 100,000 bacterial pathogens. Genome Res. 28, 1395–1404 (2018).

75. Sagulenko, P., Puller, V. & Neher, R. A. TreeTime: Maximum-likelihood phylodynamic analysis. Virus Evol. 4, vex042 (2018).

76. Seemann, T. tseemann/snippy. (2015).

77. Danecek, P. et al. Twelve years of SAMtools and BCFtools. GigaScience 10, giab008 (2021).

78. Bryant, J. M. et al. Stepwise pathogenic evolution of Mycobacterium abscessus. Science 372, eabb8699 (2021).

79. Cingolani, P. et al. A program for annotating and predicting the effects of single nucleotide polymorphisms, SnpEff: SNPs in the genome of Drosophila melanogaster strain w1118; iso-2; iso-3. Fly (Austin) 6, 80–92 (2012).

80. Abramson, J. et al. Accurate structure prediction of biomolecular interactions with AlphaFold 3. Nature 630, 493–500 (2024).

81. Sehnal, D. et al. Mol* Viewer: modern web app for 3D visualization and analysis of large biomolecular structures. Nucleic Acids Res. 49, W431–W437 (2021).

82. Jones, P. et al. InterProScan 5: genome-scale protein function classification. Bioinformatics 30, 1236–1240 (2014).

