## Supplementary material for "A 28-year evolution experiment on *Burkholderia pseudomallei* survival in nutrient-depleted sterile water"

*Indicates equal contribution

**Supplementary figure 1.** Annotated genes for putative phthalate degradation pathway in *B. pseudomallei*

**Structure of Phthalic Anhydride and hydration process**

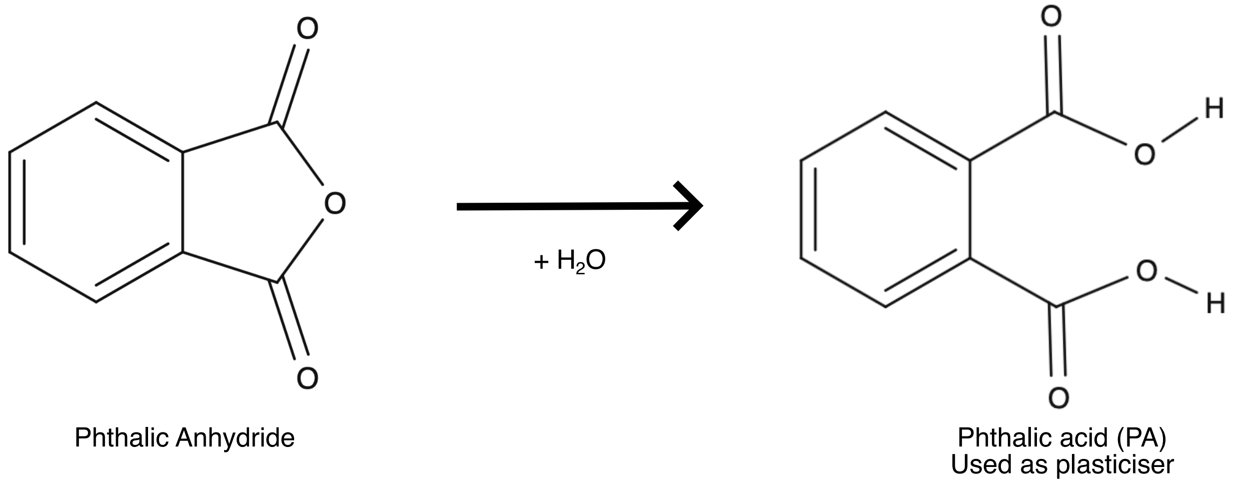

**putative *pht Operon genes***

**Protein-protein network analyses by string-db**

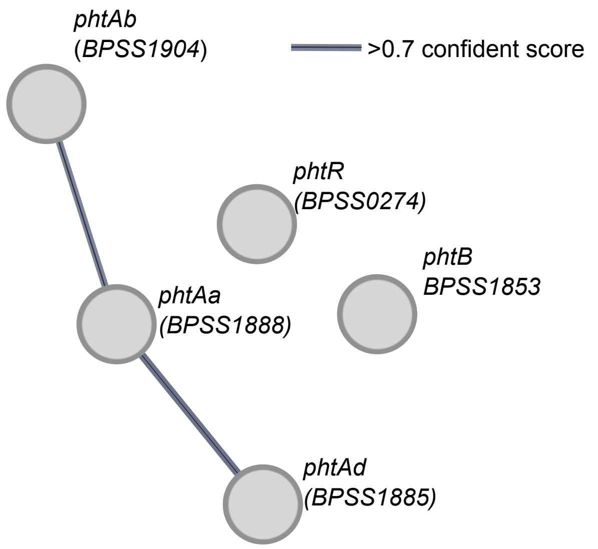

| Genes | *Mycobacterium vanbaalenii* gene (Protein id) | Homolog in *B. pseudomallei* K96243 | Amino acid seq-uence identity to K96243 (%) | Homolog in *B. pseudomallei* a207 | Gene product |
|---|---|---|---|---|---|
| *phtR* | AAQ91913.1 | *BPSS0274* | 26.78 | *A207_04309* | Pht Operon regulator |
| *phtAa* | AAQ91914.2 | *BPSS1888* | 34.88 | *A207_05335* | Phthalate dioxygenase large subunit |
| *phtAb* | AAQ91915.1 | *BPSS1904* | 31.77 | *A207_05336* | Phthalate dioxygenase, small subunit |
| *phtB* | AAQ91917.1 | *BPSS1853* | 29.21 | *A207_00570* | Dihydrodiol dehydrogena se |
| *phtAd* | AAQ91919.1 | *BPSS1885* | 36.95 | *A207_05316* | Phthalate dioxygenase, ferredoxin reductase |

**pht* operon sequence based on *M. vanbaalenii* PYR-1 accession no. AY365117. PMID: 15528661

**Supplementrary figure 2**. Number of genetic variants detected in each sample type

**a.** Number of unique single-nucleotide polymorphisms detected in each population

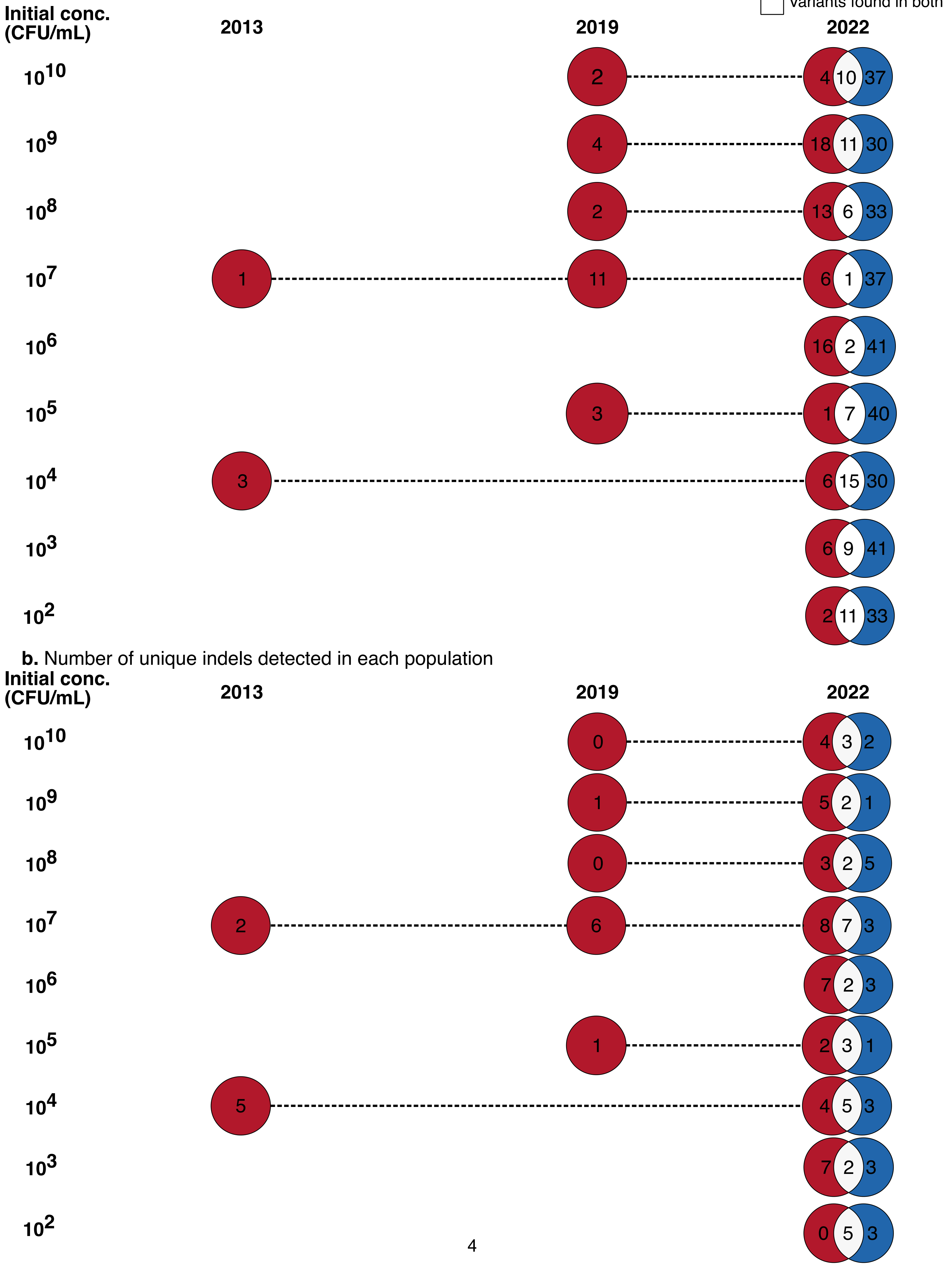

**Supplementrary figure 3**. Ancestral reconstruction of SNPs on the whole-genome phylogeny

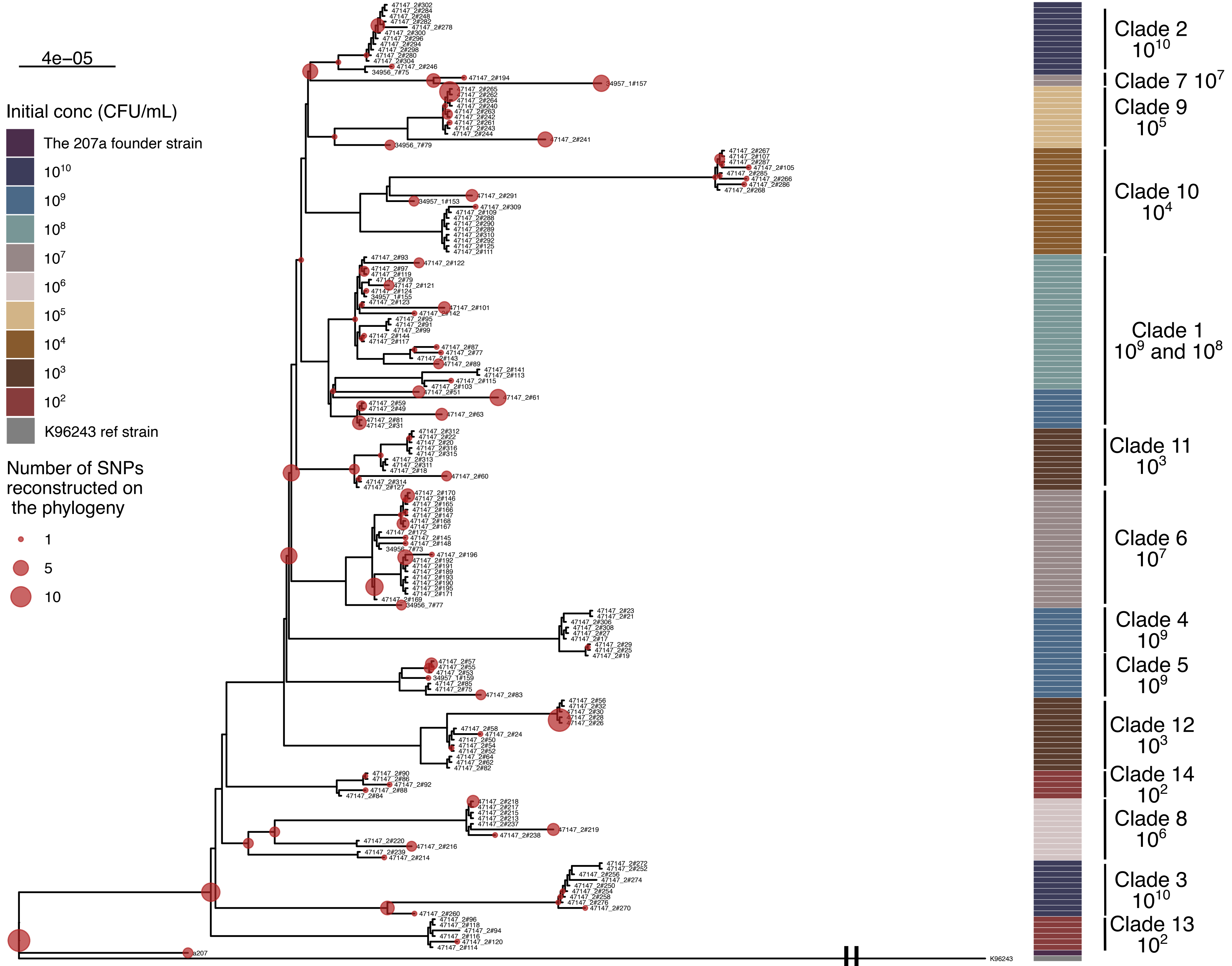

**Supplementrary figure 4**. Estimated mutational rate of the fourteen clades characterised by the ML phylogeny

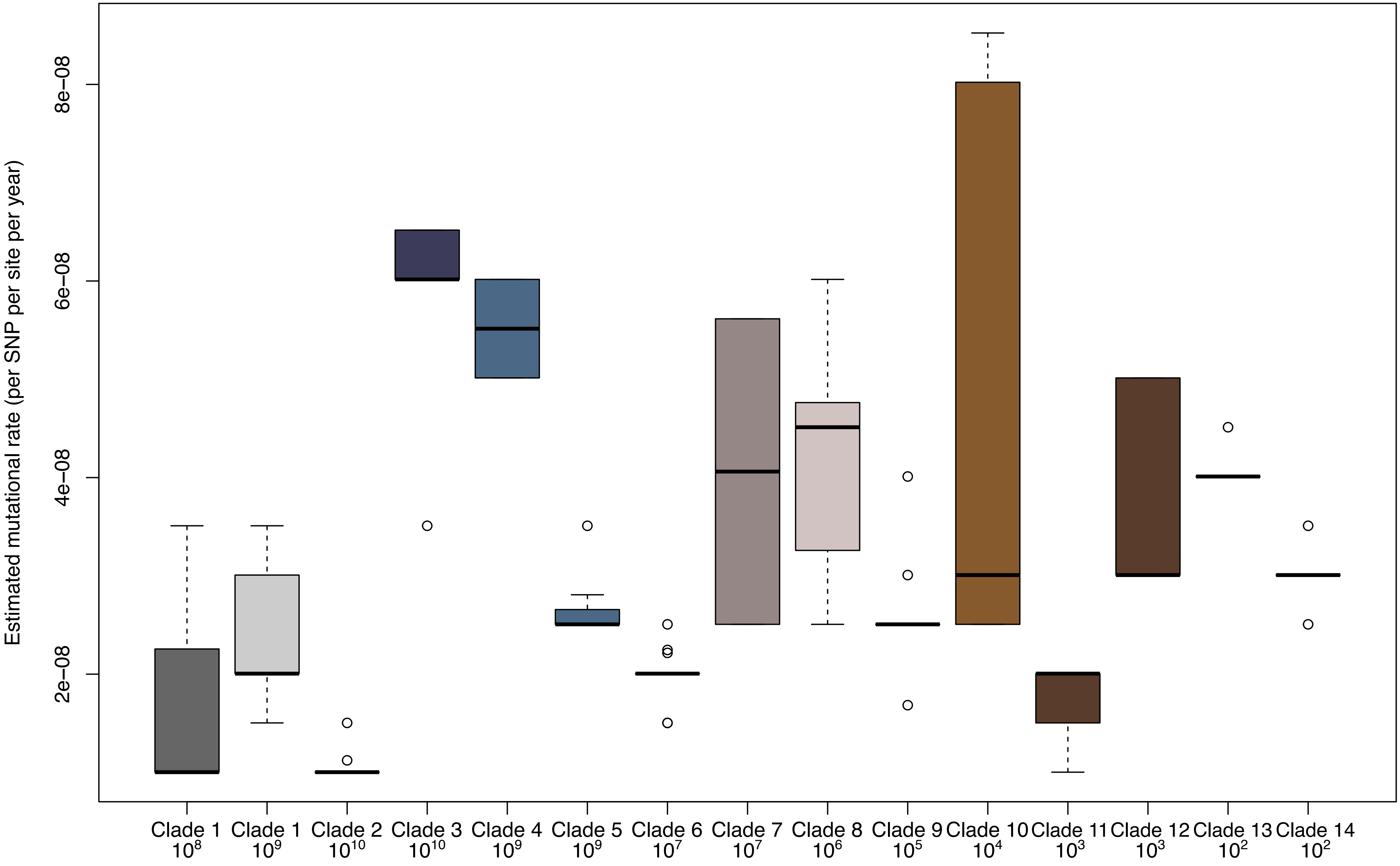

**Supplementary figure 5.** Alphafold model of *qseC* protein structure and identified domains

**a.** Alphafold model of *qseC* protein structure

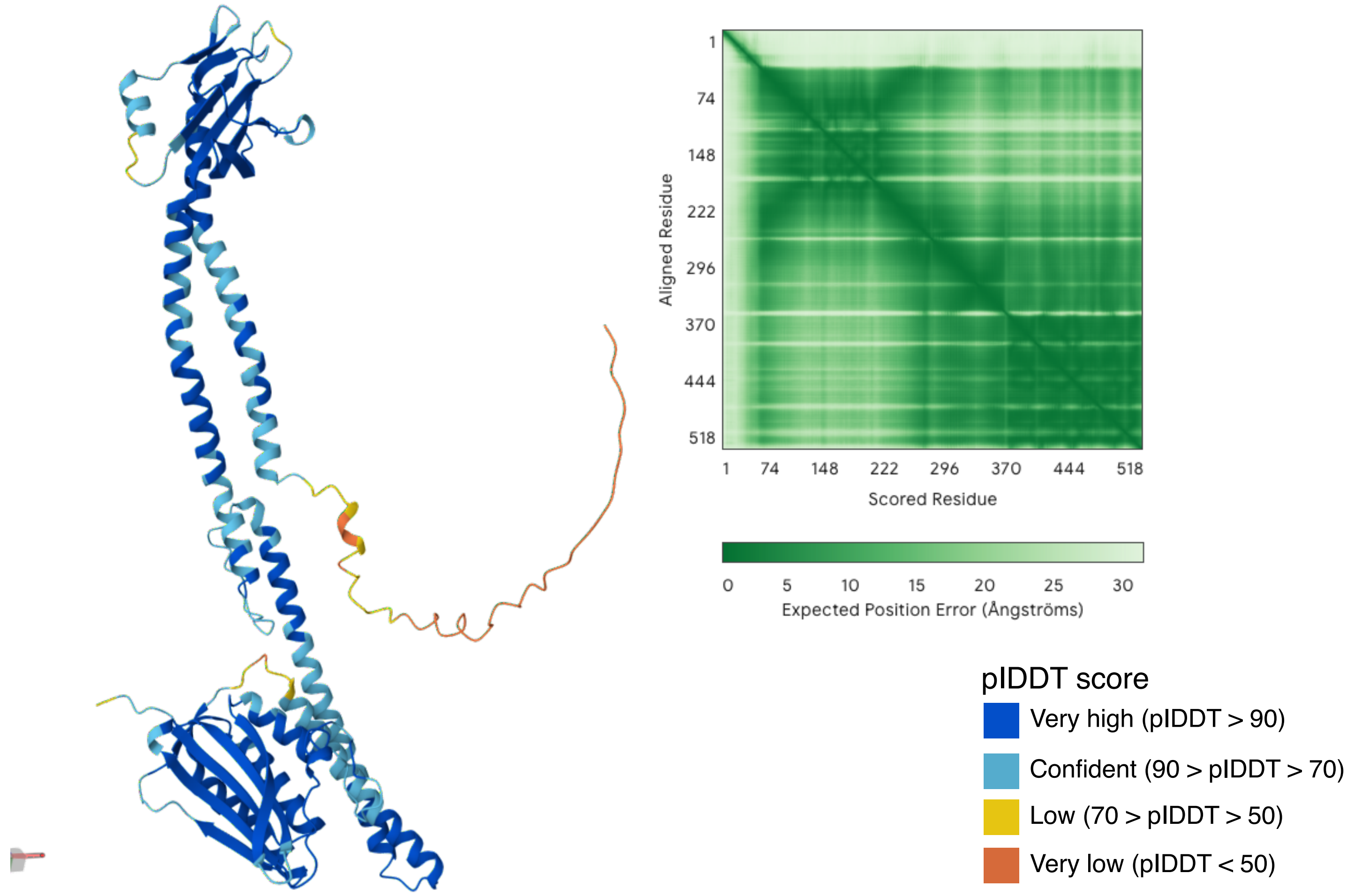

**b.** Highlighted modelled *qseC* protein with predicted domain from InterproScan

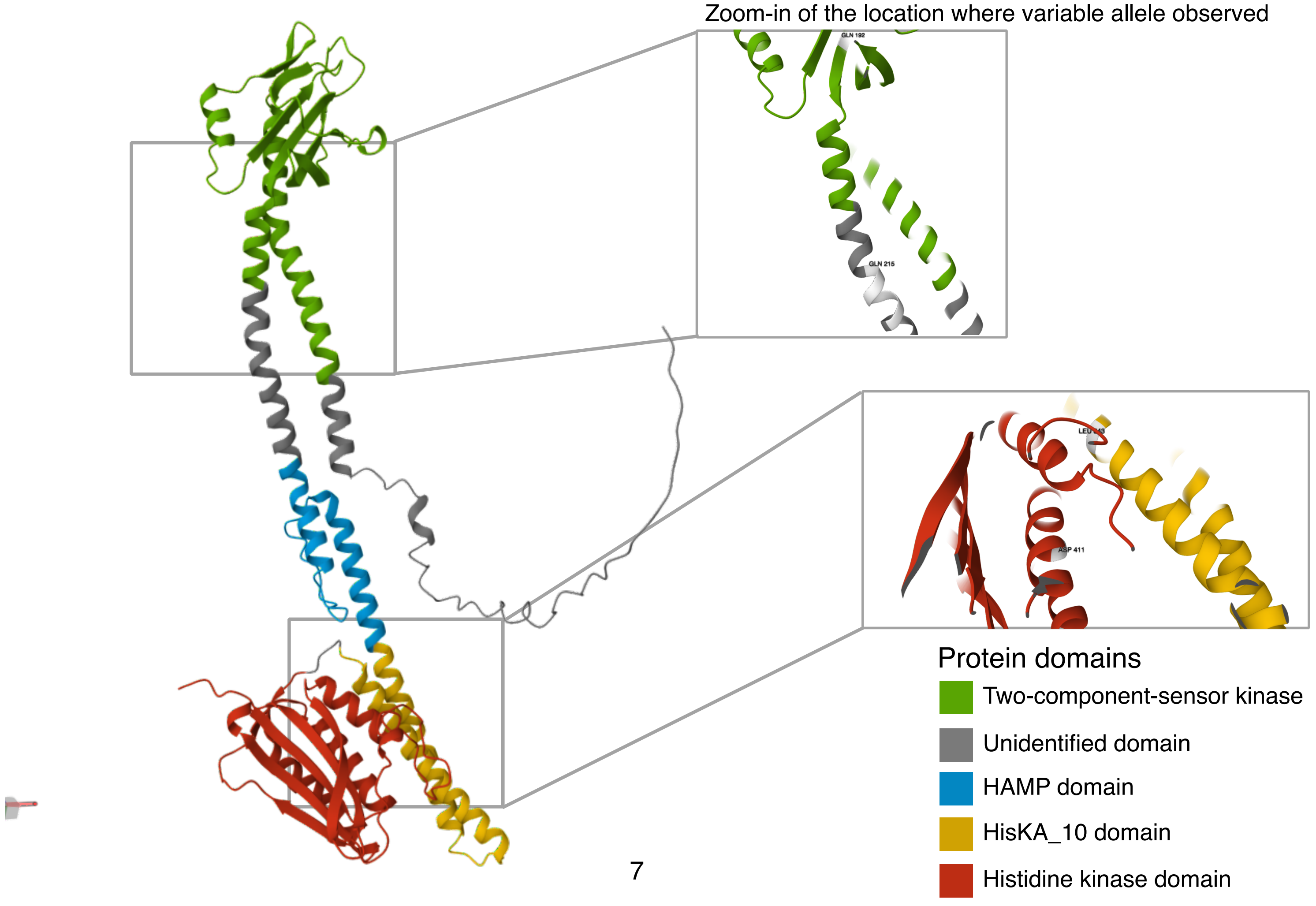

**Supplementary figure 6.** Alphafold model of *emrA* protein structure and identified domains

**a.** Alphafold model of *emrA* protein structure

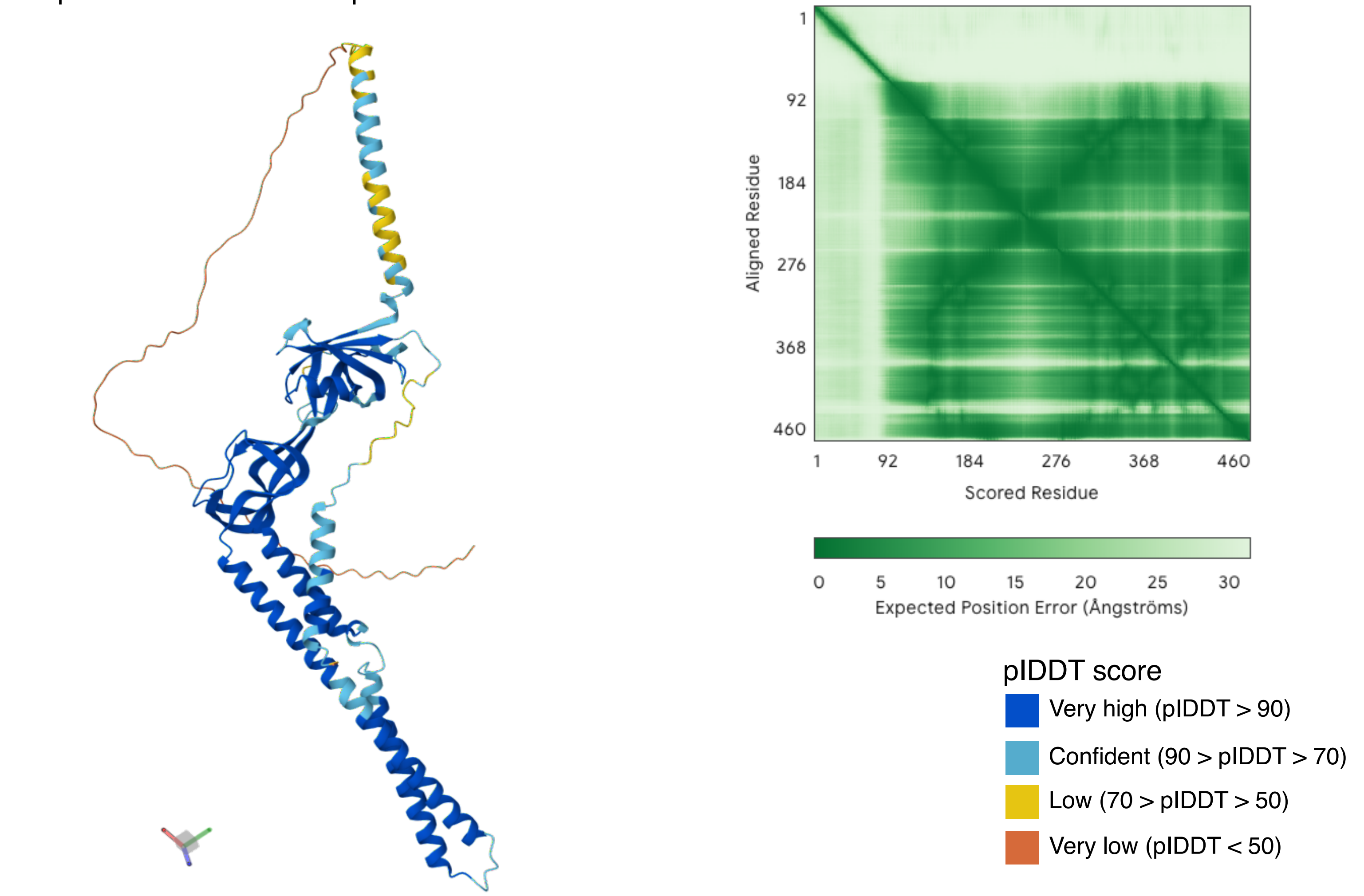

**b.** Highlighted modelled *emrA* protein with predicted domain from InterproScan

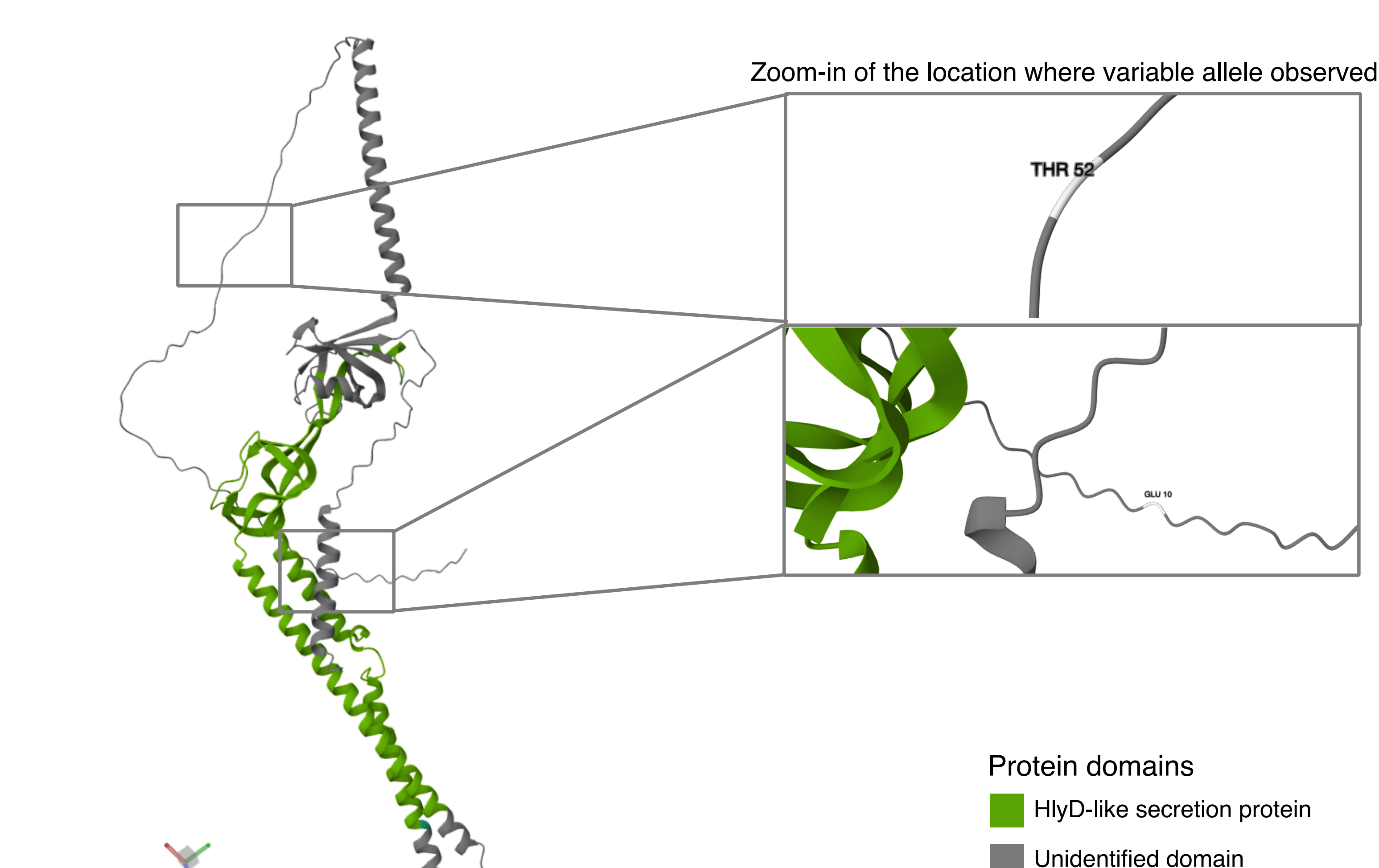

**Supplementary figure 7.** Alphafold model of *rssB* protein structure and identified domains

**a.** Alphafold model of *rssB* protein structure

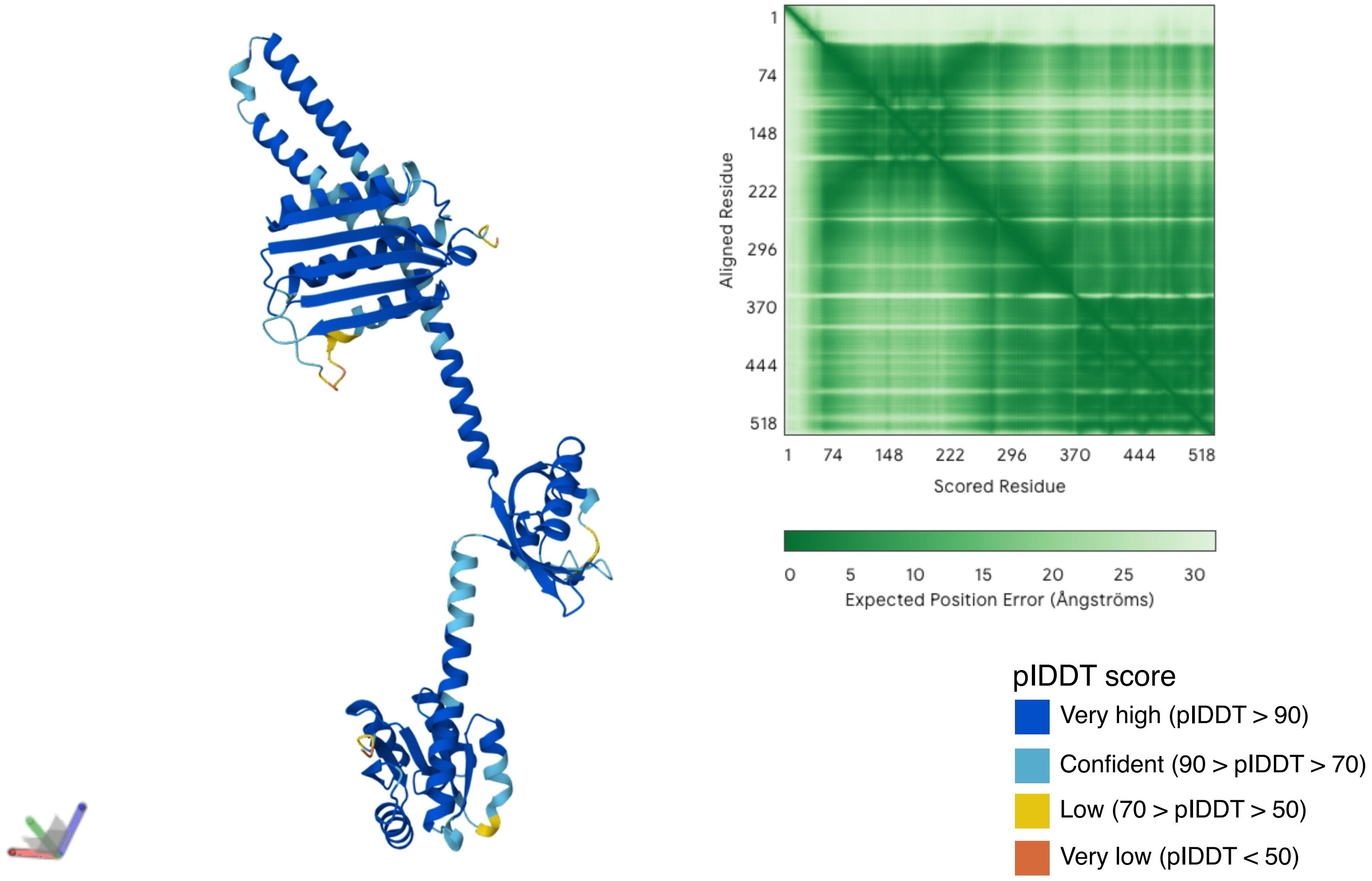

**b.** Highlighted modelled *rssB* protein with predicted domain from InterproScan

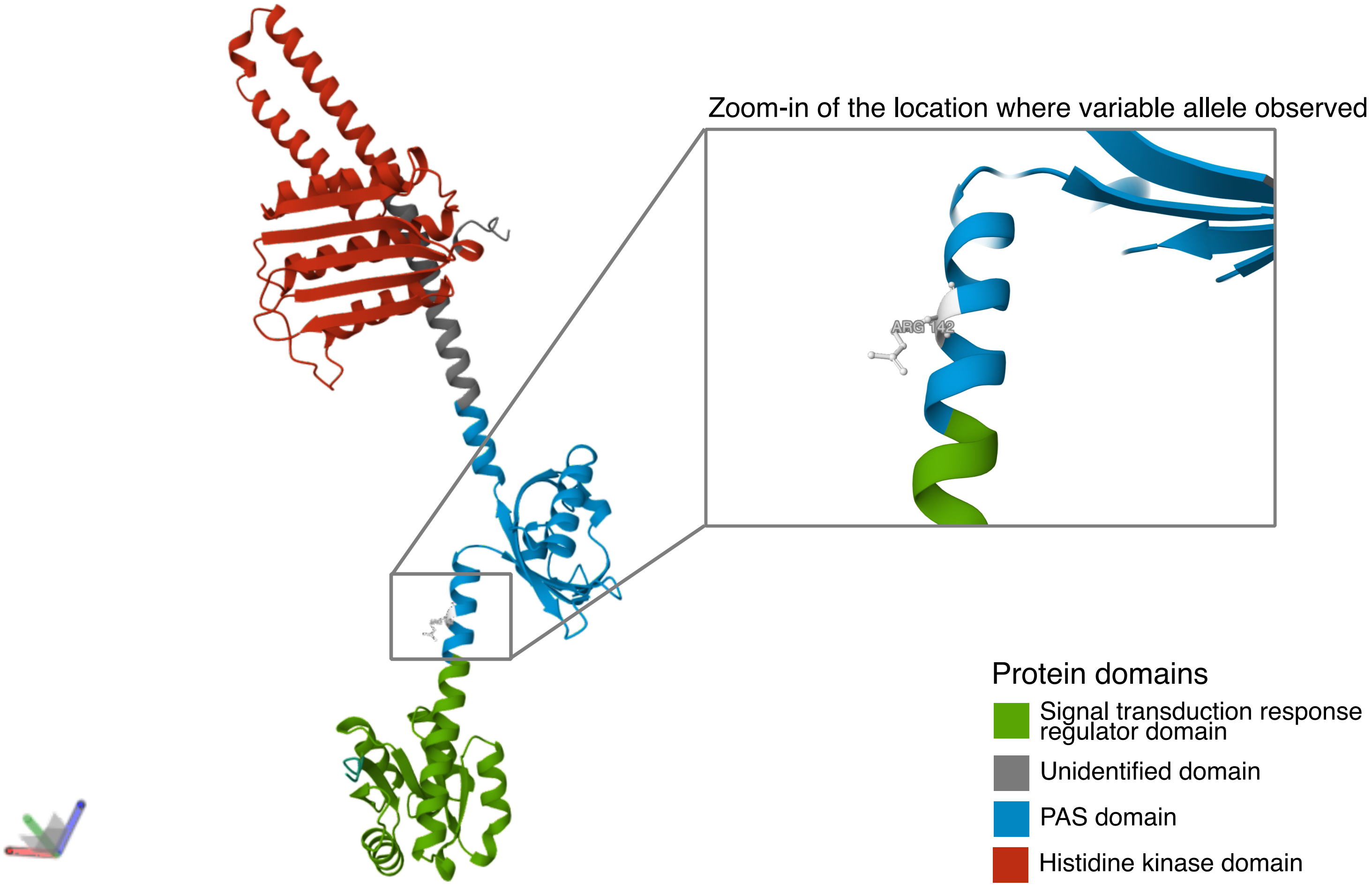

**Supplementary figure 8.** Alphafold model of *czcS* protein structure and identified domains

**a.** Alphafold model of *czcS* protein structure

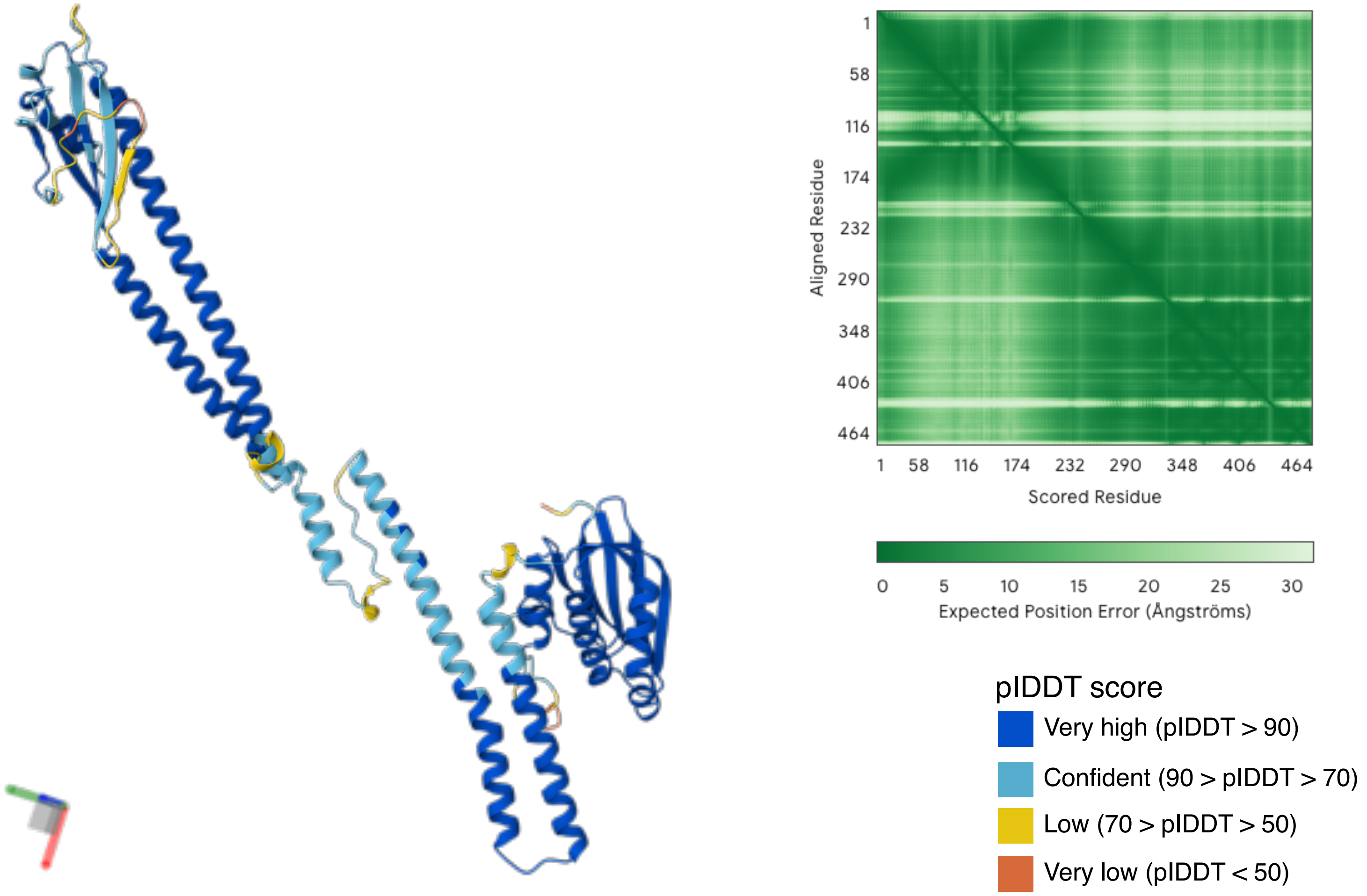

**b.** Highlighted modelled *czcS* protein with predicted domain from InterproScan

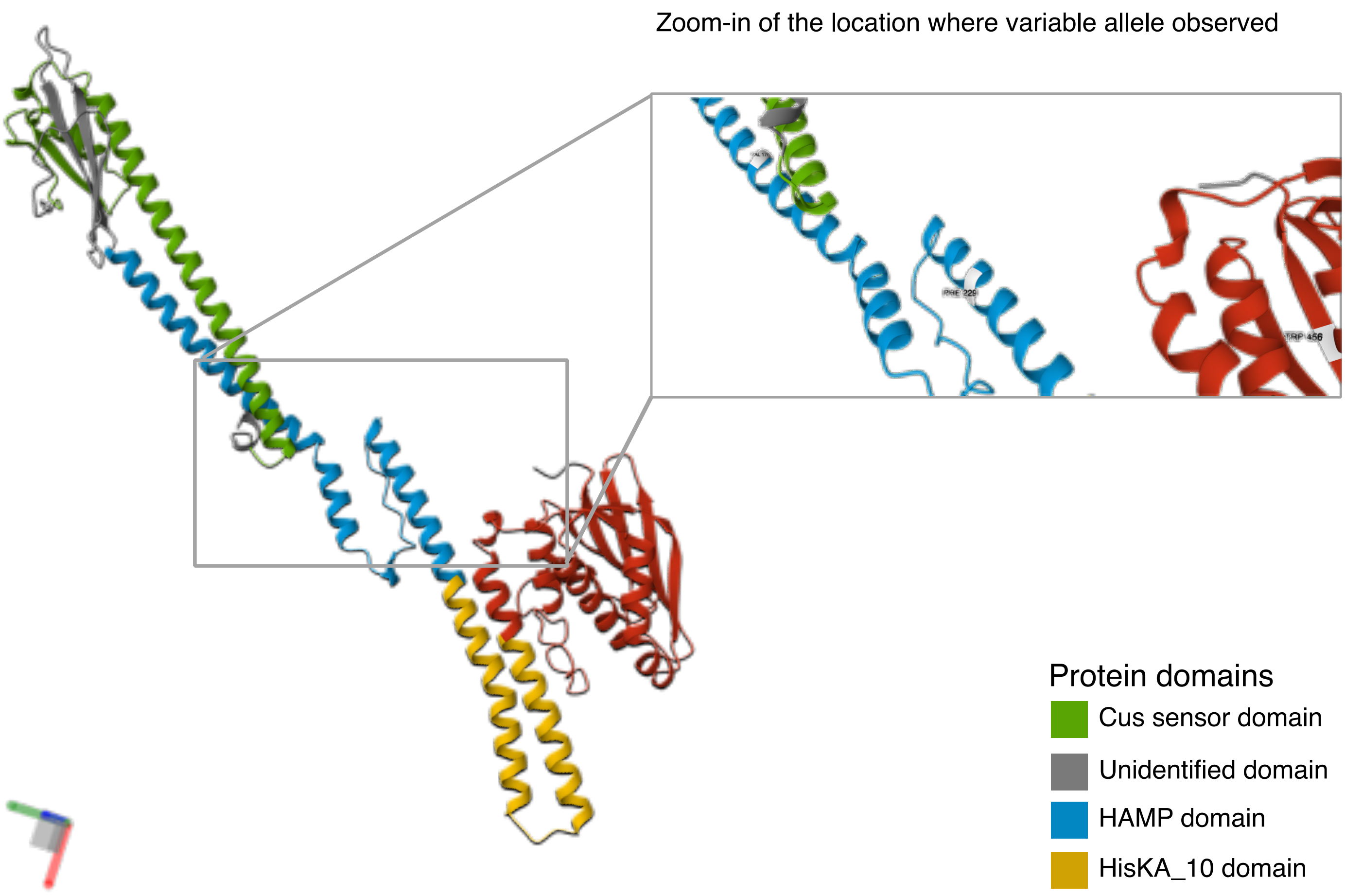

**Supplementary figure 9.** Alphafold model of *sasA* protein structure and identified domains

**a.** Alphafold model of *sasA* protein structure

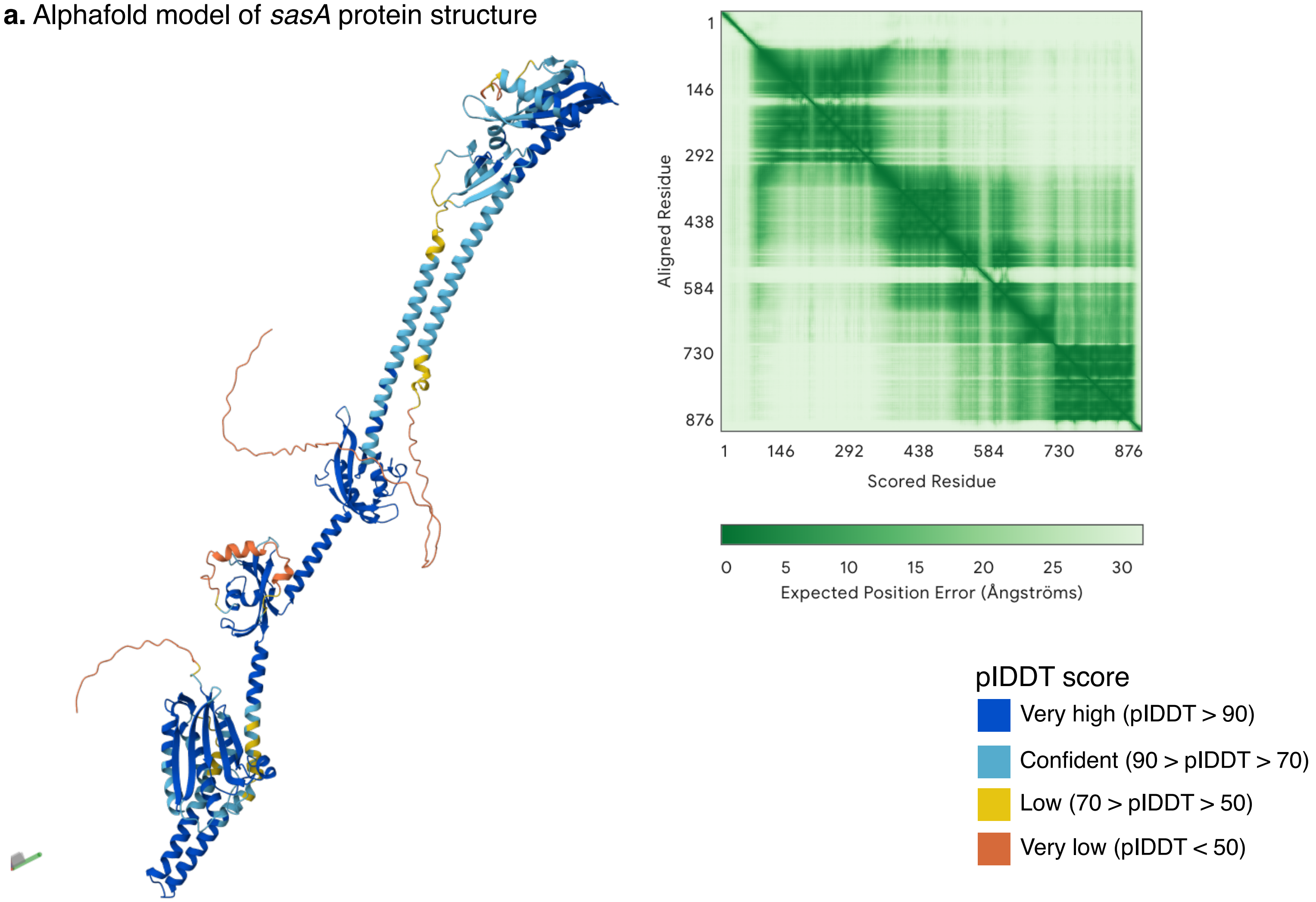

**b.** Highlighted modelled *sasA* protein with predicted domain from InterproScan

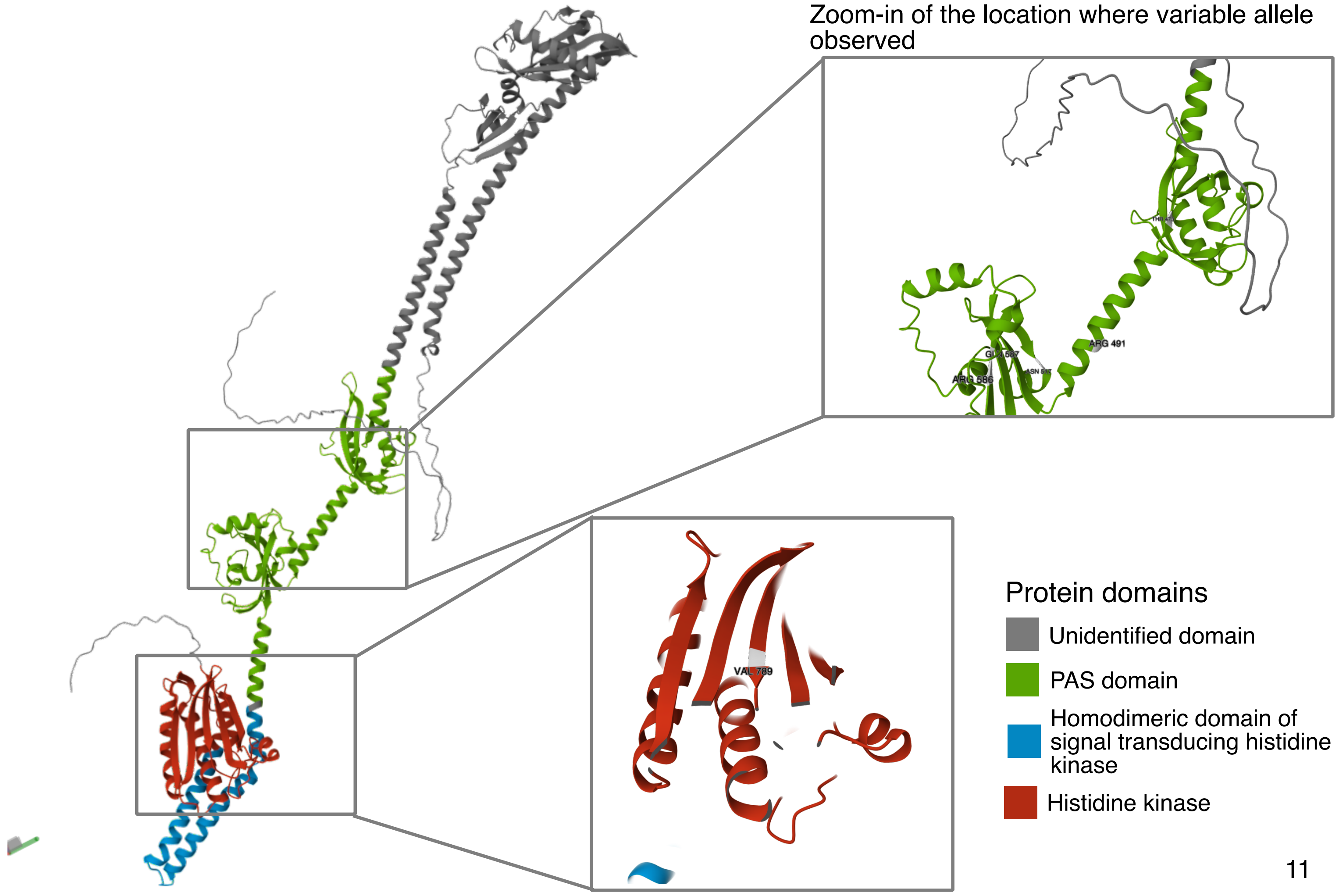

**Supplementary figure 10.** Alphafold model of *BPSL0892a* protein structure and identified domains

**a.** Alphafold model of *BPSL0892a* protein structure

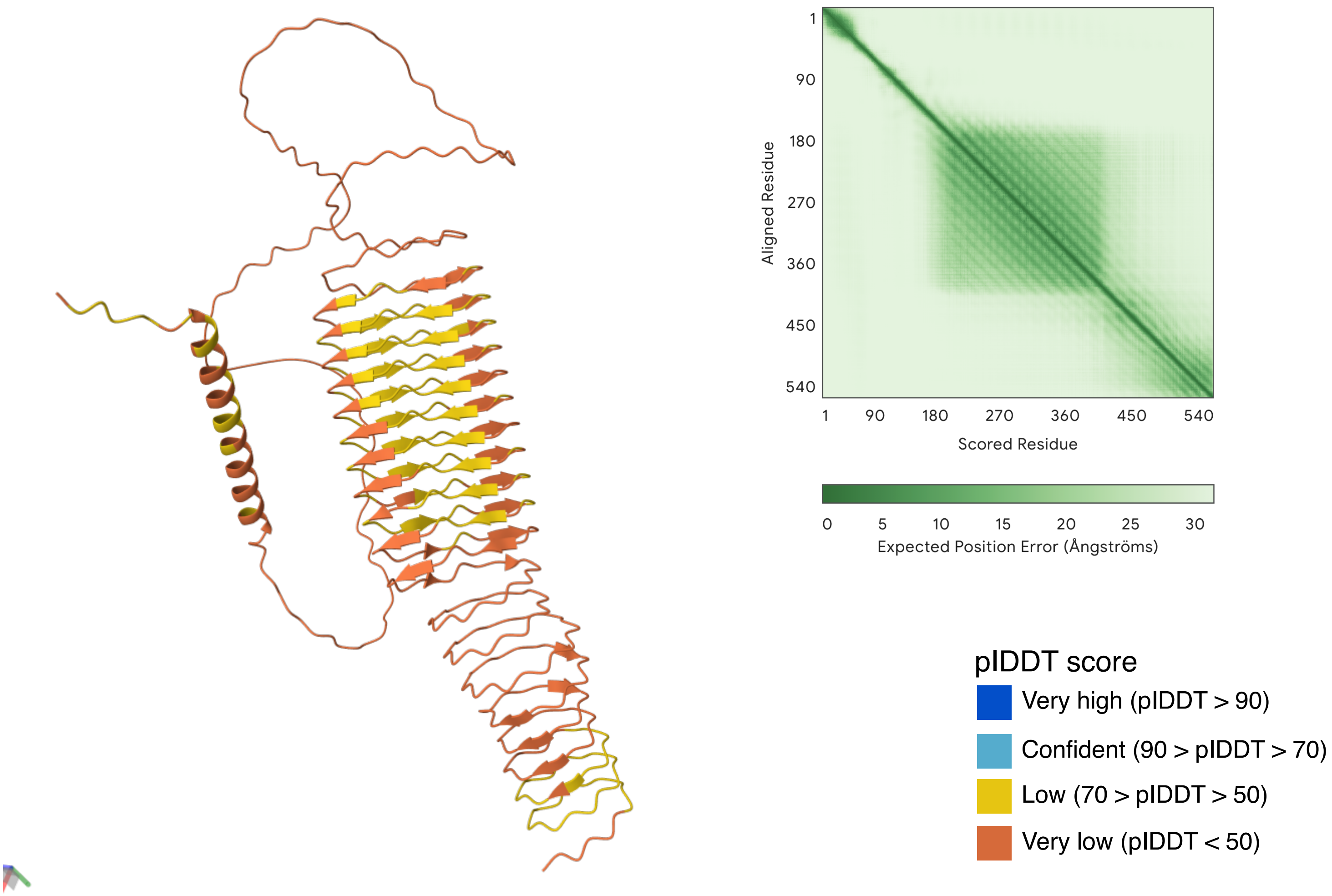

**b.** Highlighted modelled *BPSL0892a* protein with predicted domain from InterproScan

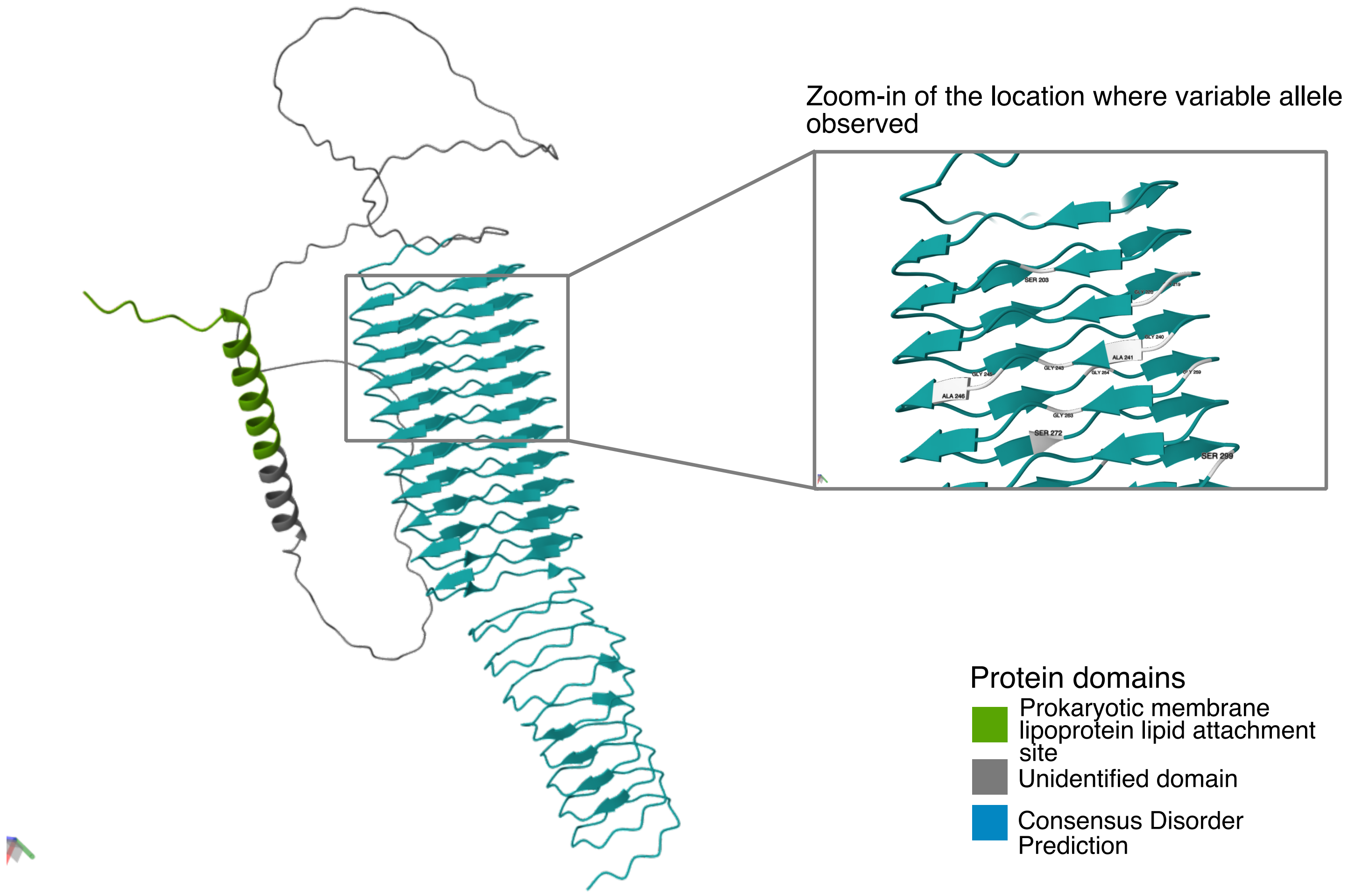

**Supplementary figure 11.** Alphafold model of *BPSS0088* protein structure and identified domains

**a.** Alphafold model of *BPSS0088* protein structure

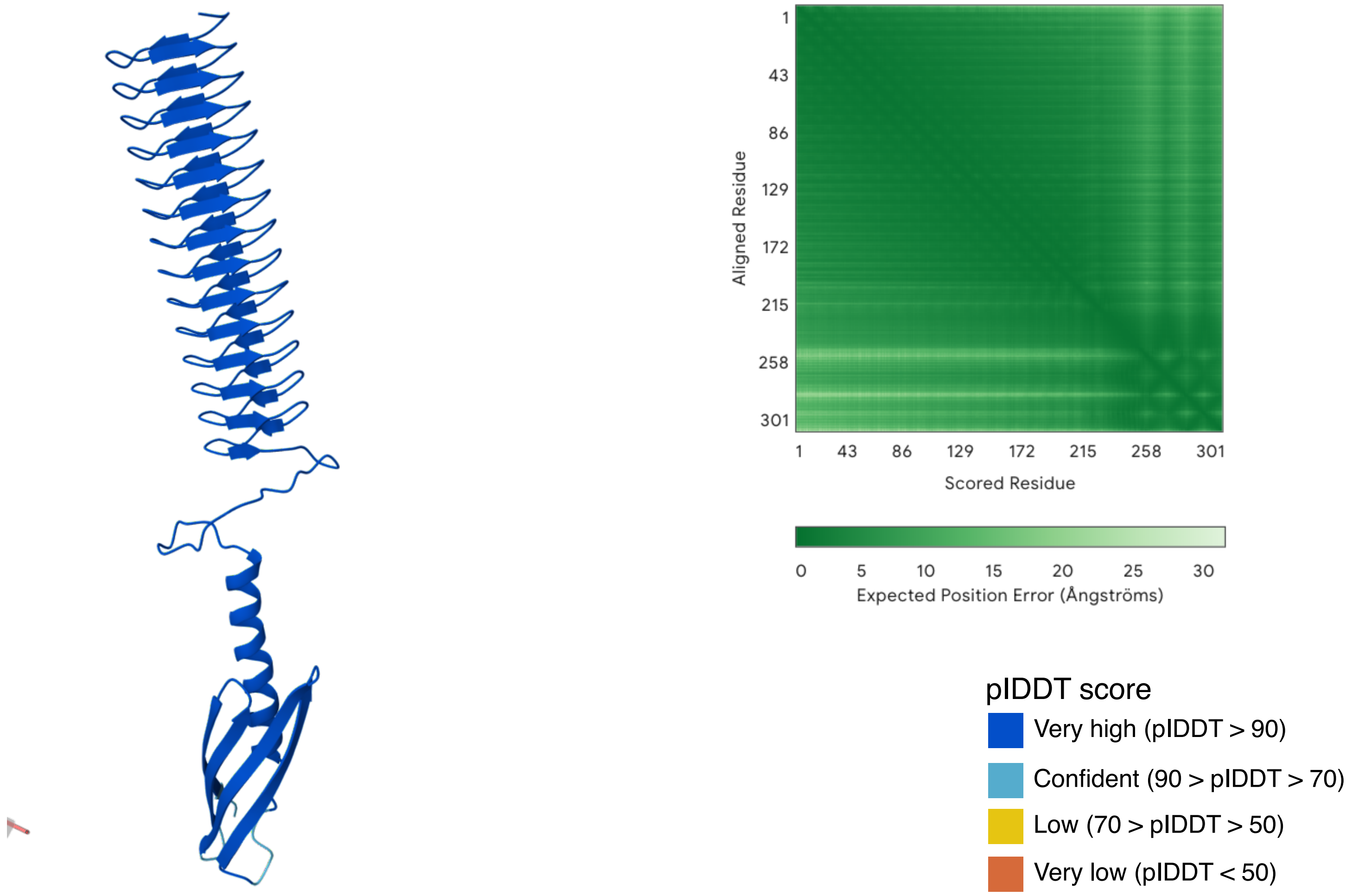

**b.** Highlighted modelled *BPSS0088* protein with predicted domain from InterproScan

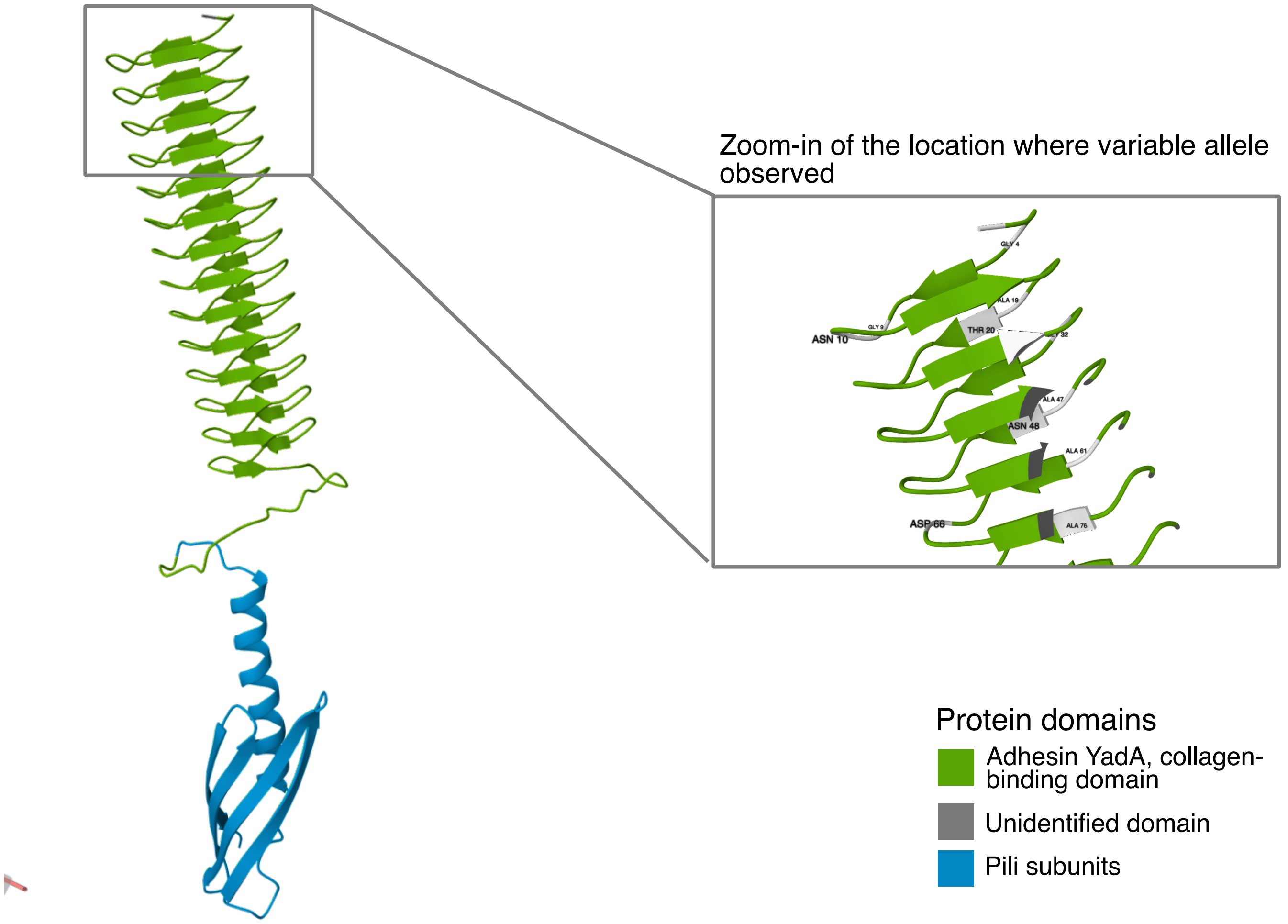

**Supplementary figure 12.** The complete genomes of the founder isolate *B. pseudomallei* 207a strain

Average GC content = 68.27 %
Predicted coding sequences = 5,787 genes

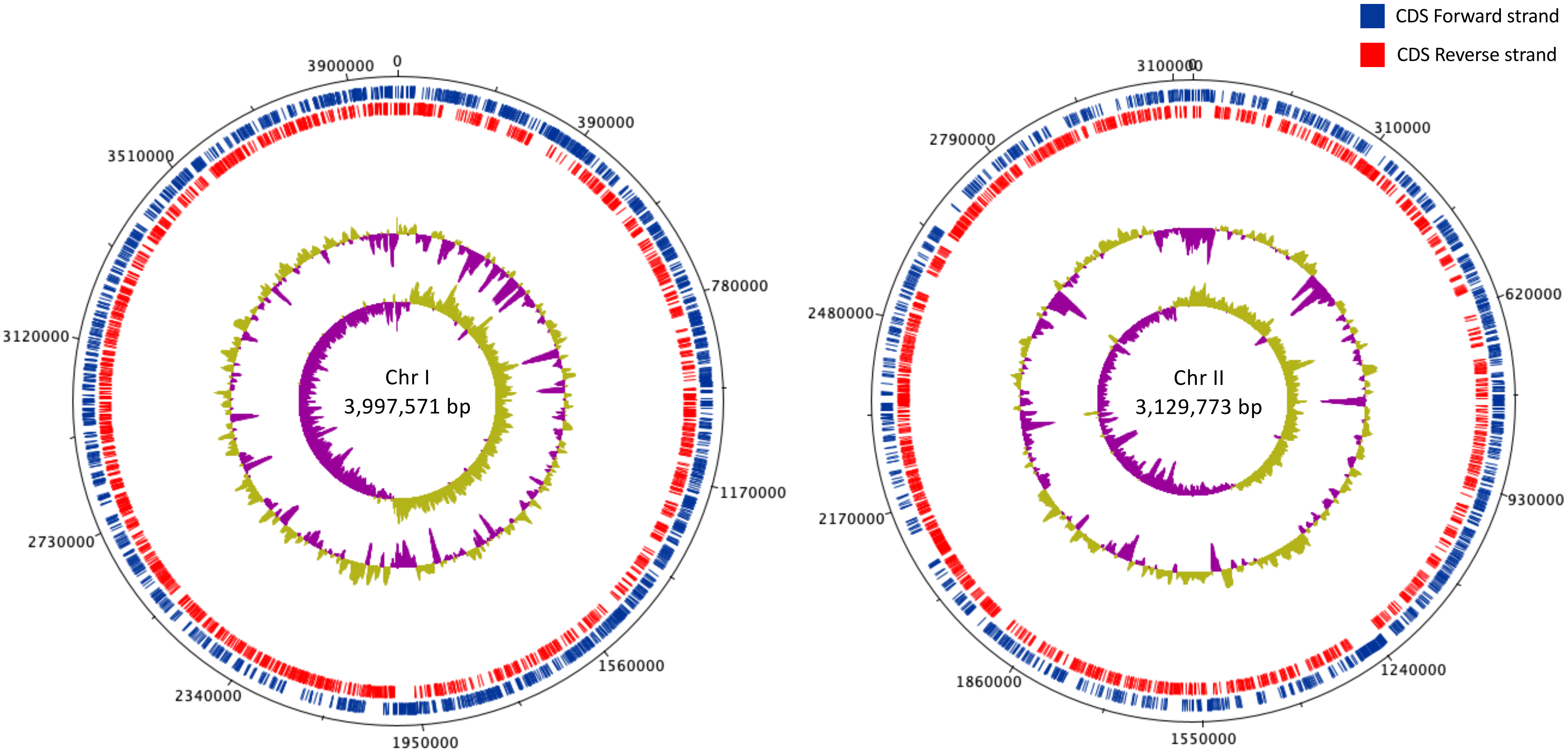
